# Molecular genetic characterization of bacterial KH-domain proteins

**DOI:** 10.64898/2026.03.16.712068

**Authors:** Kaylee T. Nguyen, Nay Won Lett, Chandra M. Gravel, Sungeun Jo, Yiling Shi, Manasa Narayan, Sahil Sharma, Cynthia M. Sharma, Katherine E Berry

**Author notes:** [Kaylee T. Nguyen] Division of Infectious Diseases, Brigham and Women’s Hospital, Boston, MA, 02115, USA; [Nay Won Lett] Department of Obstetrics and Gynecology, Brigham and Women’s Hospital, Boston, MA 02115, USA; [Sungeun Jo] Graduate Program in Molecular Medicine, University of Maryland School of Medicine, Baltimore, MD 21201, USA.

## Abstract

Post-transcriptional gene regulation is a key mechanism for bacterial stress responses and virulence, and RNA-mediated regulation frequently relies on global RNA-binding proteins (RBPs). A pair of interacting KH-domain proteins (KhpA and KhpB) have recently been identified as global RBPs in several bacterial species. To better understand their molecular functions, we employed bacterial two- and three-hybrid (B2H/B3H) assays in an *E. coli* reporter system to analyze protein-protein and protein-RNA interactions of KhpA/B orthologs from three human pathogens: *Campylobacter jejuni*, *Helicobacter pylori*, and *Clostridioides difficile*. Protein-protein interactions were conserved across all species, with KhpA-KhpB heterodimers forming more robustly than either homodimer and KhpA homodimerizing more readily than KhpB. On the other hand, protein-RNA interactions were more varied across species: *C. jejuni* and *C. difficile* KhpA bound both species-specific and non-specific RNAs, but *H. pylori* KhpA—and KhpB orthologs from all species—showed no RNA interaction in B3H assays. Site-directed mutagenesis experiments demonstrated that residues in the GXXG motif of KhpA are critical for RNA interaction and differences in these residues account for the distinct RNA-binding behaviors of KhpA orthologs. Collectively, these findings provide a cross-species, molecular view of how KhpA and KhpB recognize one another and RNA ligands to regulate gene expression.

## INTRODUCTION

Post-transcriptional gene regulation is a critical mechanism through which bacteria adapt to environmental changes, regulate stress responses, and control virulence (1–5). Central to this process are RNA-binding proteins (RBPs), which modulate the stability, translation, and decay of mRNAs through interactions with non-coding small (s)RNAs and messenger (m)RNAs. In many bacterial species, post-transcriptional regulation is mediated by global RBPs — such as Hfq and ProQ — that stabilize RNAs, recruit additional factors, and/or promote the annealing of sRNAs to target mRNAs (6–9). However, many bacterial species lack both Hfq and ProQ homologs, suggesting they may use alternative RBPs to support RNA-mediated regulation.

Recent work has identified bacterial K-homology (KH) domain proteins as a candidate class of global RBPs that are widespread across bacteria, and particularly prevalent in genomes that lack Hfq or ProQ (10). A pair of proteins containing a KH domain — KhpA and KhpB — co-occur in nearly half of bacterial genomes, including the three human pathogens examined in this study: *Clostridioides difficile*, *Campylobacter jejuni*, and *Helicobacter pylori*. Across all bacterial orthologs, both KhpA and KhpB possess an RNA-binding KH domain; KhpB proteins often include additional Jag-N and R3H domains (Figure 1), the latter of which is also an ssDNA/RNA-binding domain (11). While both KH and R3H domains typically contain conserved motifs implicated in RNA binding (GXXG and RXXXH motifs, respectively), some KhpB proteins — including those from *C. jejuni* and *H. pylori* — have degenerate/missing motifs (Figure 1A). In addition to mediating potential RNA interactions, KH domains have the ability to dimerize through an interface consisting of two alpha helices (Figure 1B; 12–14).

**Figure 1.**
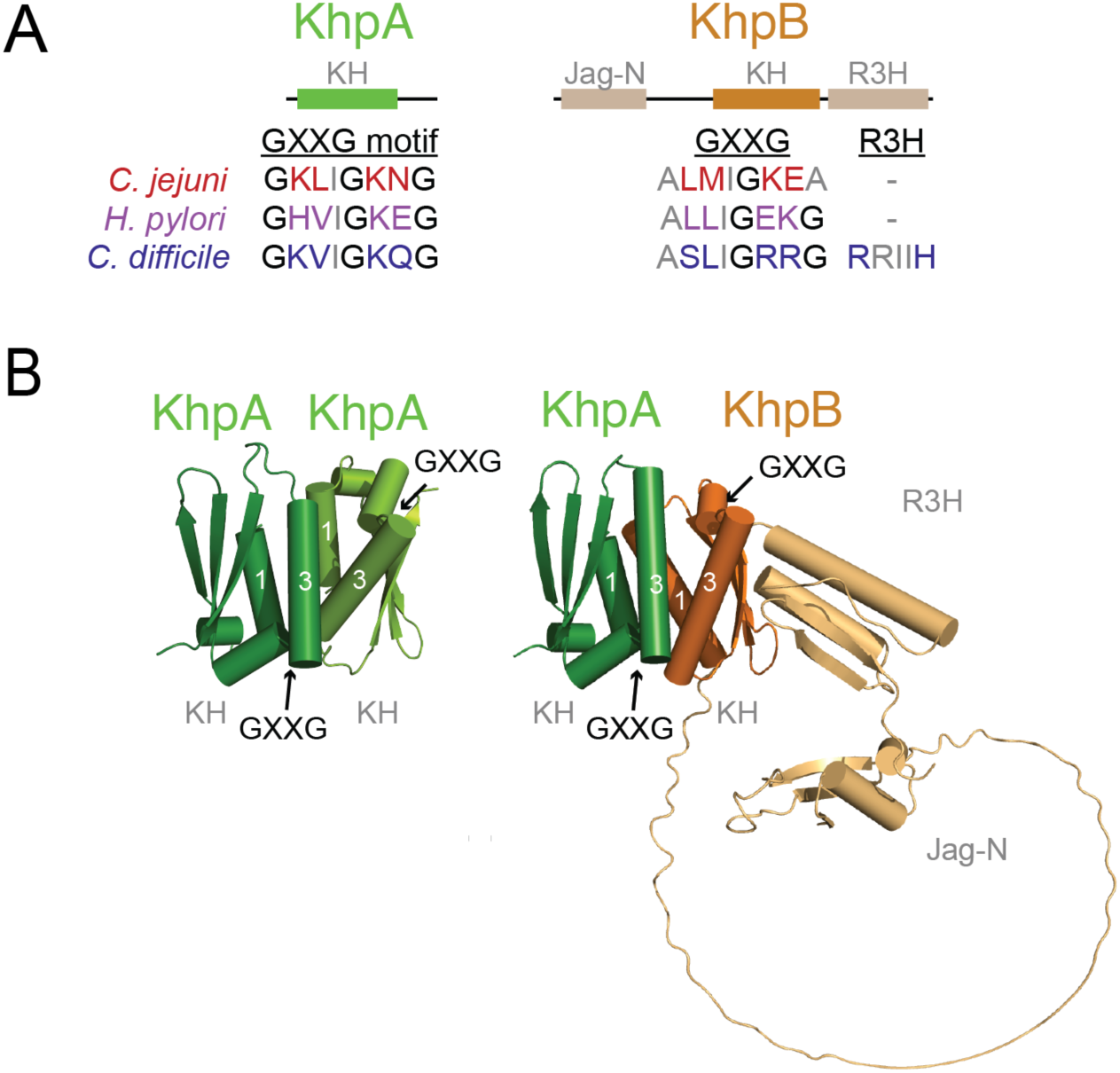
Structure of bacterial KhpA and KhpB proteins. (A) Domain structure of KhpA and KhpB. Residues within the GXXG and R3H motifs are indicated (10). Note the absence of the R3H motif residues in both *C. jejuni* and *H. pylori* KhpB as well as the degenerate GXXG motif in *C. jejuni* KhpB. (B) Predicted structure of the *C. jejuni* KhpA homodimer (left) and KhpA/KhpB heterodimer (right) showing the KH domain interface. The KH, R3H and Jag-N domains are labeled, and the locations of GXXG motif residues and predicted dimerization helices α1 and α3 are indicated. Structures were predicted using AlphaFold 3 (37).

Both KhpA and KhpB have recently been found to interact with hundred of cellular RNAs in Grad-seq and co-IP/RIP-seq experiments in both Gram-positive and Gram-negative species: *C. difficile* (15), *S. pneumoniae* (16, 17)*, Synechocystis* (18)*, E. faecalis* (19) and *F. nucleatum* (20). In addition, both proteins were recently found to co-immunoprecipate with *D. radiodurans* sRNAs (21). This broad association of KhpA/B with both sRNAs and mRNAs, coupled with the observation that deletion of either protein leads to the differential expression of many genes in several species (15, 20, 21), raises the intriguing possibility that KhpA and KhpB function as RNA-chaperone proteins, particularly in bacterial species that possess neither Hfq nor ProQ. Phenotypically, KhpA and KhpB have been implicated in cell length and division processes in *C. difficile, S. pneumoniae* and *L. planatarum* (15, 16, 22–24), as well as in the peptidoglycan stress response and virulence in *S. pneumoniae* (16, 17, 25).

In eukaryotic proteins, KH domains typically bind 4-6 nucleotides of single-stranded RNA with varying sequence preferences, and multiple KH domains within a single polypeptide can contribute combinatorially to achieve higher affinity and specificity (26). Because KhpA and KhpB each contain only a single KH domain — in contrast to eukaryotic KH proteins, which often contain multiple copies — it is possible that KhpA/B homo- and/or hetero-dimers support the recognition of longer, more complex RNA sequences than either protein alone. Consistent with this model, KhpA and KhpB have been shown to co-localize, co-sediment or co-immunoprecipitate (co-IP) in *S. pneumoniae, E. faecalis*, *Synechocystis* and *D. radiodurans* (13, 16–19, 21, 22, 27), and two-hybrid assays have detected both KhpA homodimers and KhpA/B heterodimers using *S. pneumoniae* proteins (13). However, the direct protein-protein interactions and RNA interactions of these orthologs have not been systematically compared across species.

Given the considerable variety in the amino acid sequences and electrostatic potential of KhpA/B orthologs (10), the interactions of these proteins — with themselves, one another, and RNA — may be distinct across species. With the exception of recent work in *F. nucleatum* and *D. radiodurans* (20, 21), little is known about the strength and specificity of RNA binding by isolated KhpA and KhpB proteins, or to what extent their RNA-binding behaviors are conserved across species. We chose to investigate these questions in the human pathogens *C. difficile*, *C. jejuni*, and *H. pylori*, each of which has clinical importance. While *C. difficile* is a major cause of hospital-acquired infections, the ε-proteobacteria *C. jejuni* and *H. pylori* are leading causes of gastroenteritis and gastric cancer, respectively (28–30). Because *C. jejuni* and *H. pylori* lack orthologs of global RNA-binding proteins like Hfq and ProQ (9), the Khp proteins may play an especially important role in their post-transcriptional gene regulation.

Here, we investigated the molecular interactions of KhpA and KhpB orthologs from the ε-proteobacteria *C. jejuni* and *H. pylori,* and compared them to proteins from *C. difficile*. To systematically compare these proteins, we used bacterial two- and three-hybrid assays in *E. coli.* This approach allowed us to study these orthologs in a uniform cellular environment, independent from species-specific factors, while avoiding challenges of purifying them for *in vitro* studies. Using two-hybrid assays, we found that KhpA and KhpB proteins from across all three species form robust heterodimers and that KhpA proteins — as well as the isolated KH domains of KhpB proteins — form homodimers. Interestingly, the RNA-binding activity of these proteins differed substantially in three-hybrid assays: KhpA orthologs from *C. jejuni* and *C. difficile* bound a variety of RNA sequences, while KhpB proteins and their isolated KH domains did not display binding to any RNAs tested. Mutational analysis showed that residues within the GXXG motif of KhpA contribute to RNA binding and partially account for the decreased RNA-binding activity of the *H. pylori* KhpA ortholog compared to other species. These results establish a combination of shared and unique features of bacterial KH-domain proteins across several species and set the stage for further analysis of the function of these conserved RBPs in post-transcriptional gene regulation.

## MATERIALS AND METHODS

### Biological Resources

Complete lists of plasmids, strains and oligonucleotides (oligos) used in this study are provided in Supplementary Tables S1–S3, respectively. NEB 5-α F′Iq cells (New England Biolabs) were used as the recipient strain for plasmid construction. KB473 and KB483 (*Δhfq lacZ* reporter strains (31, 32)) were used for all B2H and B3H experiments. Strains and plasmids contain antibiotic resistance genes as specified in Supplementary Tables S1 and S2, abbreviated as follows: TetR (tetracycline), KanR (kanamycin), StrR (streptomycin), AmpR (carbenicillin/ampicillin), CmR (chloramphenicol-), and SpecR (spectinomycin).

### Reagents

Restriction enzymes and additional molecular cloning reagents were purchased from New England Biolabs. Reagents specific to B2H/B3H assays were purchased from Novagen/Millipore Sigma (PopCulture Reagent, cat. no. 71092; rLysozyme, cat. no. 71110) and Gold Biotechnology (X-gal, cat. no. X4281; TPEG (phenylethyl-β-D-thiogalactopyranoside), cat. no. P-125). Materials for 96-well bacterial growth and lysis were purchased from VWR (2mL deep well blocks, cat. no. 10755–248; porous adhesive film, cat. no. 60941–086), Genesee Scientific (Olympus sterile flat bottom plates, cat. no. 25–104) and Sigma Aldrich (Greiner bio-one flat-bottom clear polystyrene plates, cat. no. 655101). Microplate shakers were purchased from VWR (cat. no. 12620–928) and Genesee Scientific (cat. no.31–213) and absorbance measurements were performed on a SpectraMax190 microplate spectrophotometer from Molecular Devices.

### Molecular cloning of B2H and B3H constructs

#### Construction of pBait and pPrey plasmids with original linkers

Plasmids for B2H/B3H experiments (pPrey-α-*Cje/Cdiff/Hpy*-KhpA/B, pBait-CI-*Cje/Cdiff/Hpy*-KhpA/B; pKN2-8, pSJ1-5, pCG110-113) were constructed by PCR amplification from genomic DNA (Supplementary Table S1) using forward and reverse primers that introduced NotI and BamHI restriction sites at the 5’ and 3’ ends, respectively. Because the *Hpy* KhpB sequence contains an internal BamHI site, NotI and BglII were used as the flanking restriction sites for constructs containing this ortholog (pCG110 and pCG111). Constructs containing isolated KhpB KH domains expressed *Cje* KhpB residues 119-198 (pKB1288) or *Hpy* KhpB residues 105-190 (pKB1289). Following PCR amplification and digestion, the resulting inserts were ligated into pKB816 (pBait-CI) or pKB817 (pPrey-α) plasmid vectors previously digested with NotI and either BamHI or BglII.

#### Construction of pBait and pPrey plasmids containing extended linker sequence

Plasmid pKN14 (pPrey-α-*Cdiff*-linker-KhpA) was constructed via mutagenic PCR from pKN2. Site-directed PCR mutagenesis was used to insert a flexible (GGGGS)_3_ linker upstream of the NotI restriction site in pKN2, using end-to-end primers (oKN16+17). The flexible linker sequence (codon-optimized via IDT) was GGTGGGGGCGGGTCTGGAGGTGGAGGTAGCGGCGGTGGTGGCAGT. Plasmid pKN15 (pBait-CI-*Cdiff*-linker-KhpA) was constructed analogously, using pKN6 as a template and primers oKN15+16. All other linker constructs were constructed by inserting the domain of interest into the NotI/BamHI sites of the parent vectors for the original pPrey-linker (pKN14) and pBait-linker (pKN15) parent vectors. PCR mutagenesis was conducted using Q5 Site-Directed Mutagenesis (New England Biolabs).

#### Construction of RNA bait constructs

The hybrid RNA plasmids containing a 5’UTR flanked by a 7-bp GC clamp (pBait-1xMS2^hp^-7GC-T*_trpA_*, pNWL2-20) were constructed by PCR amplification from genomic DNA (Supplementary Table S1) using forward and reverse primers to introduce XmaI or HindIII restriction sites at either end. Following PCR amplification and restriction enzyme digestion, these inserts were ligated into pLN43 (Nguyen et al. 2025). Plasmids pKN21-27, pNWL1 (pBait-5GC[1xMS2^hp^]) were constructed analogously and ligated into pLN28 (33),

### Beta-galactosidase assays

#### Bacterial Two-Hybrid (B2H) Assays

For liquid B2H assays, *E. coli* reporter cells (KB473 or KB483) were transformed with two plasmids: (1) a “prey” protein fused to the N-terminal domain (NTD) of the α subunit of RNA polymerase (RNAP) and (2) a “bait” protein fused to the DNA-binding λCI repressor. Negative controls were generated by replacing each hybrid component with “empty” vectors (α-empty for pPrey, and CI-empty for pBait). Following heat-shock transformation, cells were inoculated into LB medium supplemented with appropriate antibiotics (100 μg/ml carbenicillin, 100 μg/ml chloramphenicol and either 100 μg/ml kanamycin for KB473 or 100 μg/ml tetracycline for KB483). Cultures were grown overnight in 96-deep-well plates (2 ml volume) at 37°C with shaking at 900 rpm.

Overnight cultures were back-diluted (1:40) into 200 μl of fresh LB medium containing the same antibiotics and 5 μM isopropyl β-D-1-thiogalactopyranoside (IPTG) for induction of fusion proteins. Cultures were transferred to optically clear flat-bottom 96-well plates (Olympus) with sterile plastic lids and grown at 37°C with shaking (800–900 rpm) to mid-log phase (OD_600:_ 0.3–0.9). For each transformation, β-galactosidase (β-gal) activity was measured as previously described (34). Two-hybrid interactions were quantified by calculating the fold-stimulation over basal β-gal levels, defined as the β-gal activity measured in the presence of all hybrid constructs divided by the highest activity observed in negative control transformations lacking one hybrid component (pBait, pAdapter, or pPrey). Bar plots represent the mean and standard deviation (SD) of fold-stimulation values across at least three independent experiments.

#### Bacterial Three-Hybrid (B3H) Assays

Reporter cells (KB473 or KB483) were transformed with three plasmids: (1) pAdapter, expressing the CI-MS2^CP^ fusion protein (pCW17) (2) pBait, expressing a hybrid RNA composed of the MS2 hairpin (MS2^hp^) and the RNA of interest; and (3) pPrey, expressing the protein of interest tethered to the RNAP α-NTD. For negative controls, pBait and pPrey hybrid components were replaced individually with corresponding empty vectors (MS2^hp^-empty for pBait, α-empty for pPrey).

Following heat-shock transformation, cells were inoculated directly into LB medium supplemented with 0.2% arabinose (to induce RNA expression) and appropriate antibiotics (100 μg/ml carbenicillin, 100 μg/ml chloramphenicol and 100 μg/ml spectinomycin, along with either 100 μg/ml kanamycin for KB473 or 100 μg/ml tetracycline for KB483). Cultures were grown overnight, back-diluted and assayed under the same conditions described for B2H assay, except for the addition of 0.2% arabinose and 100 μg/ml spectinomycin in the back-diluted cultures. Fold-stimulation was calculated as the ratio of β-gal activity in the presence of all hybrid constructs to the highest activity observed in negative controls lacking one hybrid component. Data were averaged across at least three independent experiments, with bar plots showing the mean and standard deviation (SD).

### Statistical analyses

Data presented in all graphs represent the mean values obtained from at least three independent biological replicates. Error bars indicate the propagated standard error of the mean, calculated from the variability among replicate means. For the B2H and B3H assays examining GXXG motif variants, two-sample Student’s *t*-tests (two-tailed, assuming unequal variances) were used to assess whether the mean fold-stimulation of each variant differed significantly from the wild-type (WT) control. For B2H comparisons of constructs containing extended linkers (Supplementary Figure S2), analogous t-tests were conducted to assess all pairwise differences between constructs. Statistical analyses were performed using Microsoft Excel. *P* values < 0.05 were considered statistically significant; in figures, asterisks indicate *P* values, represented as follows: \**P* < 0.05, \*\**P* < 0.005, \*\*\**P* < 0.0005.

### Structure predictions

Secondary structure predictions of the RNA constructs used in this study were generated using RNAfold to calculate minimum free energy (MFE) secondary structures and visualized using *forna* (35, 36). Predictions of protein structures were conducted using AlphaFold3 (37).

## RESULTS

### B2H analysis of KhpA and KhpB protein-protein interactions

KH domains are known to dimerize through helices α1 and α3 (10, 12); such dimerization has the potential to form either heterodimers between KhpA and KhpB or homodimers of either protein (Figure 1B). We investigated these potential protein-protein interactions using a transcription-based B2H assay with a *lacZ* reporter (38, 39). In this system, two proteins (the “bait” and “prey”) are fused to the λCI DNA-binding protein or the N-terminal domain (NTD) of the α subunit of *E. coli* RNAP, respectively (Figure 2A). Our initial experiments utilized the original B2H constructs, each containing a triple-alanine linker between the λCI or α-NTD moiety and the proteins of interest (Figure 2B). Reporter cells bear a single-copy *lacZ* reporter downstream of a weak core promoter with an O_L_2 λ operator site centered at -62 bp relative to the transcriptional start site (Figure 2A). The O_L_2 site recruits the λCI-bait-protein fusion protein, and interaction between the bait and prey proteins stabilizes RNAP at the promoter, leading to an increase in *lacZ* transcription. An increase in β-galactosidase (β-gal) activity in the presence of both bait and prey protein compared to negative controls lacking one or the other protein therefore represents an interaction, which can be quantified as the fold-stimulation of β-gal activity relative to background levels (Figure 2C).

**Figure 2.**
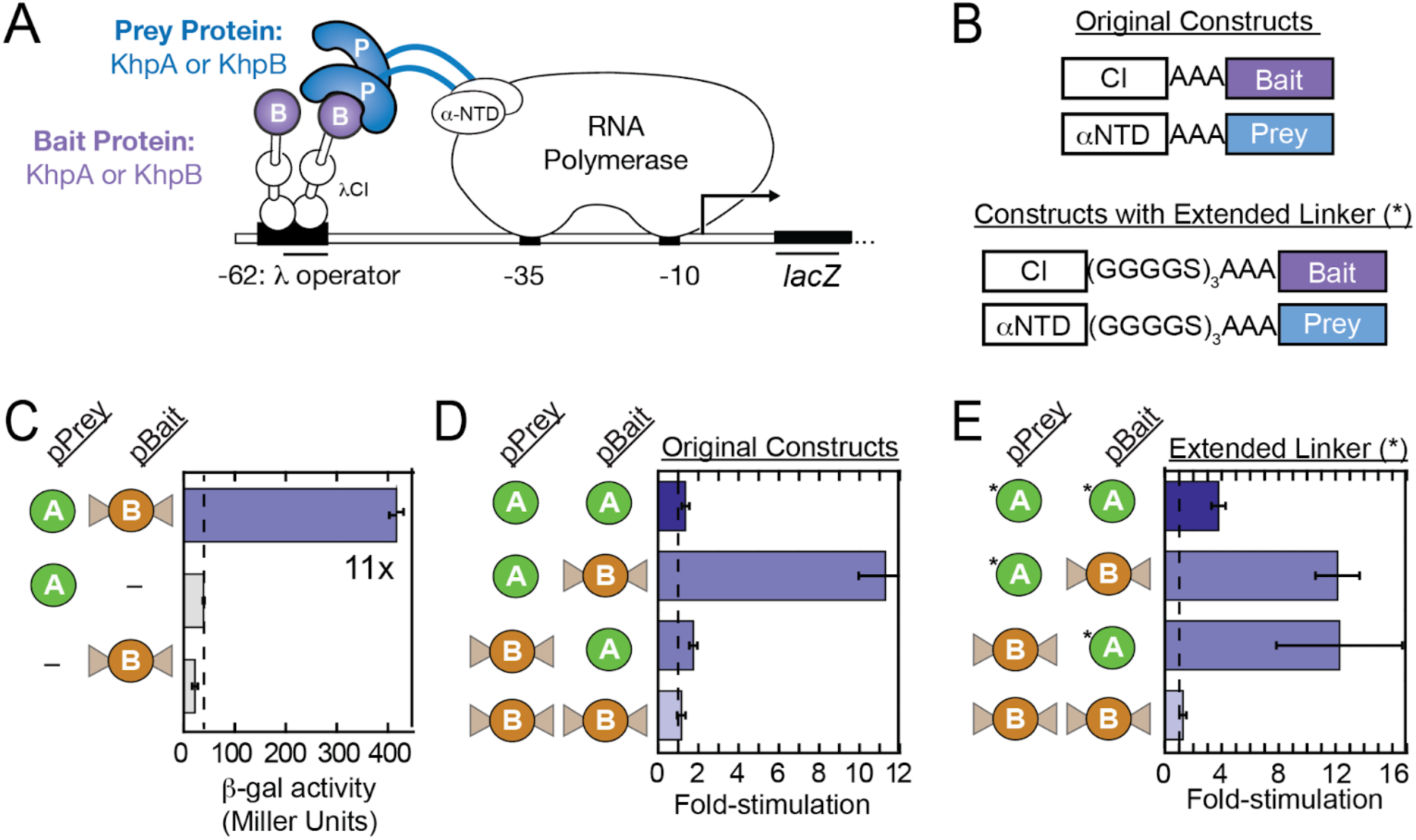
Bacterial two-hybrid assay detects KhpA-KhpB heterodimer interaction of *C. difficile* proteins. (A) Schematic of the B2H assay. Interaction between α-NTD-prey and λCI-bait fusion proteins recruit RNAP to a promoter upstream of a *lacZ* reporter gene (38). The test promoter (plac-O_L_2−62) bears the λ operator O_L_2 centered at position −62 relative to the transcription start site. Interaction between bait and prey stabilizes RNAP binding, increasing β-galactosidase (β-gal) expression. (B) Schematic of original and updated hybrid constructs. Original vectors contain a triple-alanine (AAA) linker between the λCI or α-NTD moieties and the protein of interest. Extended constructs introduce an additional flexible linker (GGGGS)₃, denoted by an asterisk within figure schematics throughout the paper. (C) B2H assay detecting heterodimerization of *Cdiff* KhpA and KhpB. Reporter cells were transformed with pPrey encoding α alone (–) or the α-KhpA fusion protein (pKN2), and pBait encoding λCI alone (–) or CI-KhpB (pKN8). In this and all subsequent panels, bar graphs represent mean β-gal activity from three biological replicates with error bars indicating standard deviation (SD). The dashed line denotes basal β-gal levels used to calculate fold-stimulation. (D) B2H assay detecting homo- and heterodimerization using combinations of *Cdiff* KhpA (pKN2, pKN6) and KhpB (pKN4, pKN8). Bar graphs in this panel and throughout the paper present B2H interactions as fold-stimulation over basal levels. (E) Effect of extended linkers on the detection of *Cdiff* B2H interactions using combinations of *Cdiff* KhpA with extended linkers (pKN14, pKN15) and KhpB (pKN4, pKN8).

We began our analysis using the *Clostridioides difficile (Cdiff)* proteins in the original B2H constructs. Interaction between *Cdiff* KhpA and *Cdiff* KhpB proteins produced an ∼11-fold stimulation of *lacZ* activity over background levels when fused to α-NTD (pPrey-KhpA) and λCI (pBait-KhpB), respectively (Figure 2C). We next sought to compare heterodimerization of KhpA and KhpB with homodimerization of each protein. While each of these fusion proteins interacted with its heterodimeric partner, neither KhpA nor KhpB homodimers were detected with these constructs (Figure 2D). Notably, KhpA/B heterodimerization appeared much weaker when the proteins were presented in the opposite orientation (pPrey-KhpB and pBait-KhpA), leading to only ∼2-fold stimulation.

The striking asymmetry of heterodimer detection between the two orientations of bait and prey proteins led us to consider whether one or both of the fusion proteins might be geometrically constrained within the transcription complex. While KhpB has a long linker preceding its KH domain (Figure 1), the KH domain in KhpA lies at the immediate N-terminus of the protein. Given that one of the predicted dimerization helices of KhpA is located at the protein’s immediate N-terminus, we hypothesized that the dimerization interface of KhpA might be constrained by its fusion to the α-NTD or λCI moieties. We therefore created constructs that introduced an extended, flexible 15-amino-acid linker sequence — (GGGGS)_3_ — beyond the triple-alanine linker present in original pBait and pPrey constructs (Figure 2B; 32, 38). Because the Jag-N domain and linker region of KhpB provide a natural linker to either α-NTD or λCI fused to its N-terminus, we only introduced the extended linkers into the KhpA constructs.

When B2H analysis was conducted again with both pBait- and pPrey-KhpA fusions containing the extended (GGGGS)_3_ linker, the *Cdiff* KhpA/B heterodimer was observed in both orientations (Figure 2E), consistent with the possibility that one of the original KhpA constructs had been geometrically constrained. Interestingly, using these extended-linker constructs, KhpA homodimerization was also detected, although at a reduced level relative to KhpA/B heterodimers (∼4- vs. ∼12-fold stimulation of β-gal activity).

### KhpA/B heterodimerization and KhpA homodimerization is conserved across multiple bacterial species

Having established that heterodimerization of the *C. difficile* KhpA and KhpB proteins is more robustly detected than homodimerization of either protein, and that extended linkers support the detection of KhpA homodimers, we next sought to compare this behavior to orthologs from *Campylobacter jejuni* (*Cje*) and *Helicobacter pylori* (*Hpy*). The *Cje* and *Hpy* KhpA and KhpB proteins share 37% and 42% amino acid identity, respectively, whereas the *Cdiff* orthologs are more divergent, sharing ∼21–23% (KhpA) and ∼19–20% (KhpB) identity with the other two species (Supplementary Figure S1). B2H analysis using constructs containing extended linkers revealed that heterodimerization is generally preferred over homodimerization across all three species, with heterodimerization exhibiting up to 4-fold and 7-fold stimulation of *lacZ* activity for *Cje* and *Hpy* proteins, respectively (Figure 3). In addition, homodimerization of KhpA could be detected for both *Cje* and *Hpy* proteins, though with slightly lower fold-stimulation values than those observed for the *Cdiff* proteins. In contrast, KhpB homodimerization was not detected for any of the species tested.

**Figure 3.**
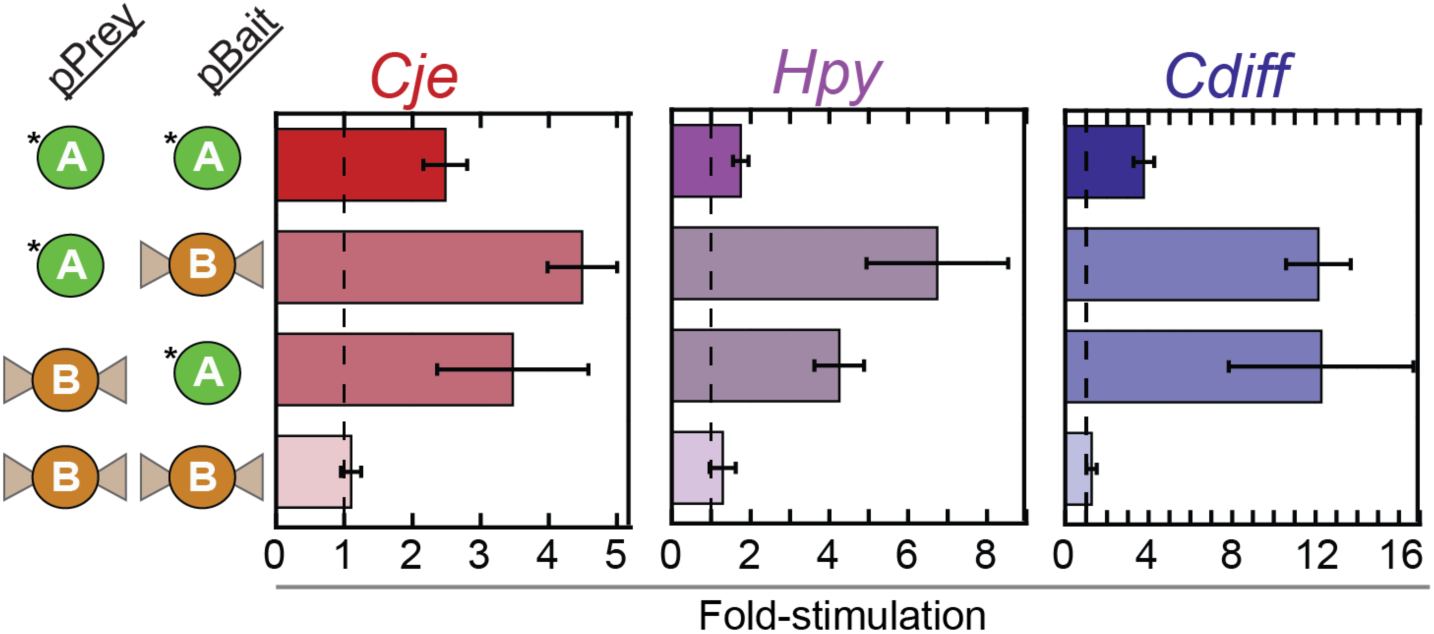
Conservation of KhpA/B heterodimerization and KhpA homodimerization. B2H results testing homo- and heterodimerization of KhpA and KhpB proteins from *C. jejuni* (red; pKN16, pKN29, pCG112, pCG113), *H. pylori* (purple; pKN17, pKN30, pCG110, pCG111) and *C. difficile* (blue; pKN4, pKN8, pKN14, pKN15). Transformations included combinations of these plasmids or corresponding empty vectors to establish basal transcription levels. Asterisks indicate the use of the extended flexible linker in KhpA constructs. Note that *C. diff* data are replotted from Figure 2E for comparison.

Given that homodimerization of *Cdiff* KhpA was only detectable using KhpA constructs with extended linkers (Figure 2D vs. 2E), we sought to determine the extent to which extended-linker constructs supported the detection of *Cje* and *Hpy* protein-protein interactions in Figure 3. To address this, we systematically compared B2H interactions across all three species by introducing the extended linker to KhpA constructs on one or both sides of the interaction. Introducing the extended linker on either the pPrey or pBait construct improved detection of *Cje* KhpA homodimerization (Supplementary Figure S2), with an additional increase observed when both constructs possessed the (GGGGS)_3_ linker. For the *Cdiff* KhpA interaction, the extended linker on the pBait(λCI) construct caused a dramatic increase in signal, with only a modest additional increase when the linker was present on both constructs. In contrast, detection of *Hpy* KhpA homodimerization was not affected by the extended linkers on either construct.

With respect to heterodimer interactions, the extended linker on the pBait construct improved the signal for both *Hpy* and *Cdiff* proteins and led to more symmetric results between the two possible orientations of the proteins (bait vs. prey). On the other hand, the linker on the pPrey construct had a greater positive effect on the *Cje* heterodimer interaction. Overall, although the magnitude of the effect varied between species, the extended linker did not reduce the strength of any interaction examined. Together, these comparisons indicate that the extended linker — particularly on the pBait(λCI) constructs — improves the detection of protein-protein interactions of KhpA orthologs.

To further explore the specificity of the KhpA-KhpB heterodimer interaction, we next asked if the KhpA and KhpB proteins of *C. jejuni* and *H. pylori* — sharing 25% and 39% sequence identity, respectively — could engage in cross-species interactions. Interestingly, while *Cje* KhpA interacted robustly with *Hpy* KhpB, no interaction between *Hpy* KhpA and *Cje* KhpB was detected (Supplementary Figure S3A). This was true regardless of whether the KhpA and KhpB proteins were fused to the pBait or pPrey construct. As a control, both *Hpy* KhpA and *Cje* KhpB hybrid proteins that failed to interact with each other nevertheless displayed strong interactions with their cognate partners from the same species (Figure 3 and Supplementary Figure S3B). AlphaFold3 predictions suggest a structural basis for this asymmetry: the “failed” *Hpy* KhpA/*Cje* KhpB pairing brings two positively charged lysine residues into close proximity (Supplementary Figure S3C), creating electrostatic and steric hindrance that likely prevents the interaction. The species-specificity of the heterodimer interactions serves as a control for the specificity of the B2H interactions and suggests that the dimerization interfaces of these KH domains have evolved to be distinct between species.

### The Jag-N and R3H domains of KhpB limit its dimerization

Given that KhpB homodimerization was not detected in the B2H assay for any ortholog tested and that dimerization is predicted to occur through the KH domain (Figure 1B), we wondered whether the additional Jag-N and R3H domains of KhpB might interfere with the ability of its KH domain to participate in protein-protein interactions. To test this possibility, we cloned the isolated KH domains into pPrey and pBait constructs containing the extended linkers described above. Although we were unable to obtain functional constructs for the isolated KH domain of *Cdiff* KhpB, we successfully examined the interactions of the isolated KH domains from the *Cje* and *Hpy* KhpB orthologs.

We first compared the heterodimer interactions between species-matched KhpA proteins and either full-length (FL) KhpB proteins or the isolated KH domains (Supplementary Figure S4A). Strikingly, the isolated KH domain of KhpB demonstrated a much stronger interaction with KhpA than the FL KhpB protein for both species. We next examined the homodimerization of KhpB using a combination of FL constructs and their isolated KH domains. For both the *Cje* and *Hpy* proteins, the isolated KhpB-KH domain showed negligible interactions with FL KhpB in either orientation (Supplementary Figure S4B). However, isolated KhpB-KH domains from each species did display a modest homodimerization interaction (∼2-fold). These fold-stimulation values were similar to those observed for KhpA homodimerization of the *Cje* and *Hpy* proteins, but lower than those of *Cdiff* KhpA (Figure 3). Overall, these results suggest that, at least in the context of this B2H assay, the Jag-N and/or R3H domains of KhpB reduce the dimerization activity of the KhpB KH domain.

#### KhpA orthologs from *C. jejuni* and *C. difficile* bind RNA in a B3H assay

Having characterized the protein-protein interactions of KhpA and KhpB in the B2H assay, we next investigated the RNA-binding behavior of each protein using the closely related B3H assay, which has previously been used to detect and characterize RNA interactions of the RNA-binding proteins Hfq and ProQ (31, 32, 34). The B3H method uses the same pPrey construct as the B2H system and functions analogously, but with the addition of a λCI-MS2^CP^ fusion protein that bridges the -62-O_L_2 site to a hybrid RNA consisting of a 5′ MS2 hairpin (MS2^hp^) upstream of a bait RNA of interest (Figure 4A).

**Figure 4.**
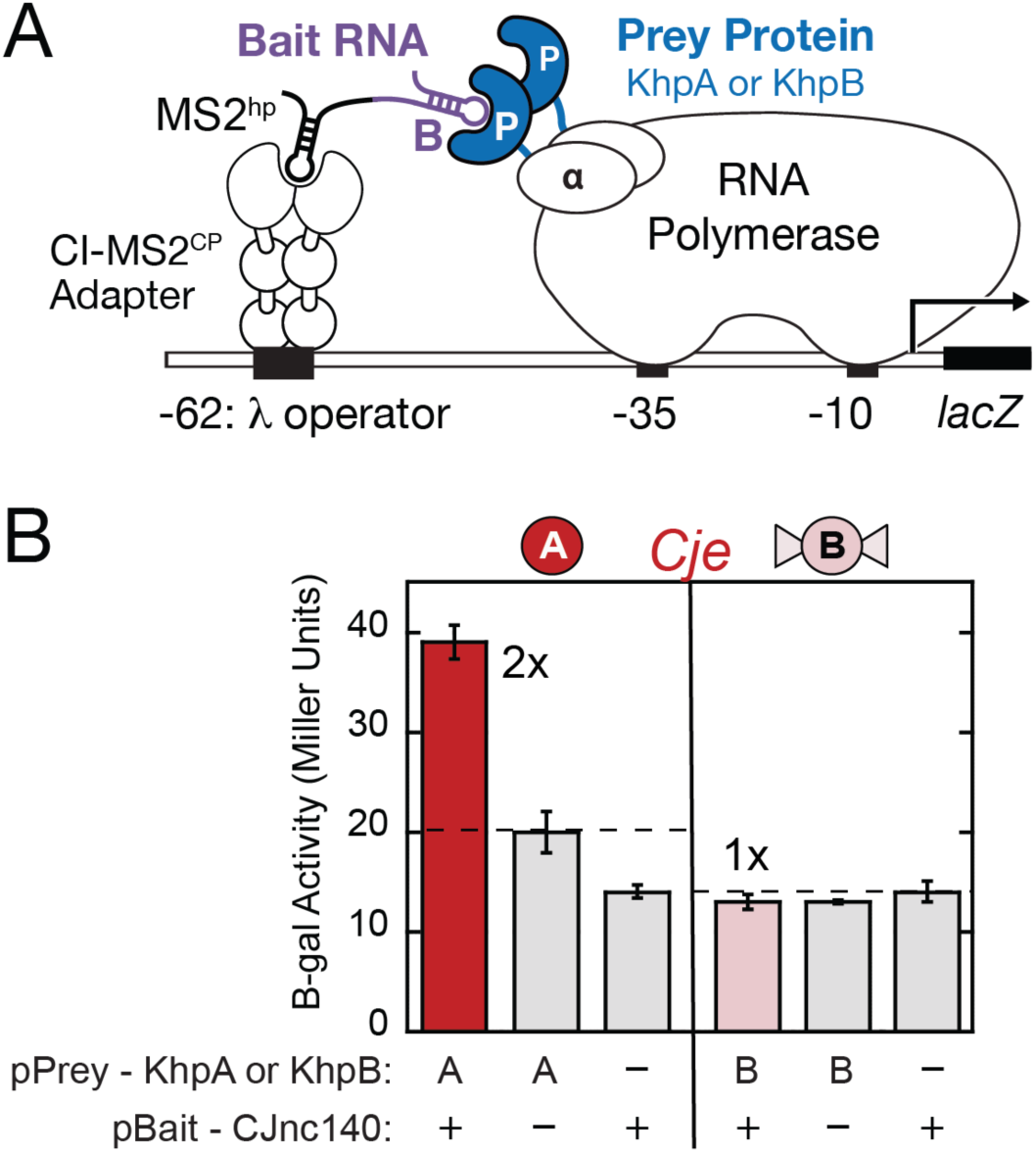
B3H-based detection of *C. jejuni* KhpA-RNA interaction. (A) Schematic of the B3H system. Interaction between an α-NTD-prey fusion protein and an MS2 hairpin (MS2^hp^)-RNA bait moiety activates *lacZ* expression from the same test promoter used in B2H assays (Figure 2); the CI–MS2^CP^ tethers the hybrid RNA bait to the promoter. (B) Results of β-gal assays. Reporter cells (KB483) were transformed with three compatible plasmids: pPrey encoding α alone (–), or KhpA (pSJ1) or KhpB (pCG112) fusions; pAdapter encoding the CI–MS2^CP^ fusion protein (pCW17); and pBait encoding a hybrid RNA containing CJnc140 flanked by a 5-bp GC clamp (pNWL1), or a negative control RNA that contained only the MS2^hp^ moiety and GC clamp (–, pSS1).

In the B3H system, these components are encoded on the pAdapter (λCI-MS2^CP^) and pBait-RNA (MS2^hp^-RNA) plasmids, respectively. Quantification of protein-RNA interactions follows the same logic as for B2H interactions: the fold-stimulation of *lacZ* is calculated as the β-gal activity in the presence of all three hybrid components, normalized to the highest value of the negative controls missing either the prey protein or bait RNA (Figure 4B). We chose to use an *Δhfq E. coli* reporter strain to avoid the potential for endogenous *E. coli* Hfq to compete with the heterologously expressed KH proteins and because the *Δhfq* strain has been shown to improve signal for both Hfq- and ProQ-RNA interactions (31, 32, 40).

We began by examining the B3H interaction of *Cje* KhpA and KhpB with the small non-coding sRNA CJnc140 (41), which had been enriched in KhpB RIP-seq experiments (Sharma lab unpublished data; details will be reported elsewhere). When both the KhpA prey and the CJnc140 bait RNA were present in B3H experiments, roughly two-fold higher β-gal activity was detected relative to negative controls (Figure 4B). This stimulation of β-gal activity indicates an interaction between *Cje* KhpA and the CJnc140 sRNA. In contrast to this association between KhpA and the sRNA, the pPrey-*Cje*KhpB construct yielded no stimulation over background levels.

To explore interactions with a broader range of RNAs, we selected a panel of RNAs from *C. jejuni, H. pylori* and *C. difficile* to serve as B3H bait RNAs. These candidates were identified based on reports of their *in vivo* association with one or both of the Khp proteins through RIP-seq experiments (*Cdiff* (15); *Cje and Hpy*, Sharma lab unpublished data, details will be reported elsewhere). These included seven RNAs from *C. jejuni* (one ncRNA and six mRNA fragments); five mRNA fragments from *H. pylori;* and six ncRNAs from *C. difficile.* In addition, we included additional *E. coli* bait RNAs previously used to detect interactions with Hfq and ProQ (31–33) and a control construct with two tandem MS2 hairpins (2xMS2^hp^). All other pBait constructs contained only a single MS2^hp^, to avoid the potential interaction with a second MS2 hairpin, as was previously observed for *E. coli* ProQ (31)). Non-coding RNAs that contain an intrinsic terminator sequence were cloned into a pBait construct downstream of the MS2^hp^, while 5′UTRs and ORF regions from mRNAs were cloned into a pBait construct in between the MS2^hp^ and an exogenous intrinsic terminator (T*_trpA_*) at the 3′ end (33). Secondary structure predictions of bait RNAs are shown in Supplementary Figures S5-S8.

The binding of KhpA and KhpB proteins from *Cje, Hpy* and *Cdiff* was assessed in the B3H assay against panels of 5-7 species-specific RNA baits and 2-5 control *E. coli (Ec)* RNAs. Robust RNA interactions with several RNAs were observed for KhpA proteins from *Cje* and *Cdiff*, but not for *Hpy* KhpA (Figure 5; raw β-gal data showing specific negative controls for each pBait construct are shown in Supplementary Figure S9). Both *Cje* and *Cdiff* KhpA interacted with approximately half of the tested RNAs. They also showed strong interaction with at least one *Ec* RNA as well as the construct containing two tandem MS2^hp^ structures, suggesting that KhpA is capable of non-specific interactions with RNAs from other species. In contrast, no RNA interaction was detected for any of the three KhpB prey proteins tested (Figure 5; pale bars). Importantly, all pPrey fusion proteins that failed to interact with RNAs in the B3H assay (i.e., *Hpy* KhpA and *Cje, Hpy and Cdiff* KhpB proteins) demonstrated robust heterodimerization activity in our B2H experiments (Figure 3). This B2H control indicates that, although these proteins do not interact with RNA in the B3H assay, they are stably expressed and correctly folded inside the cell.

**Figure 5.**
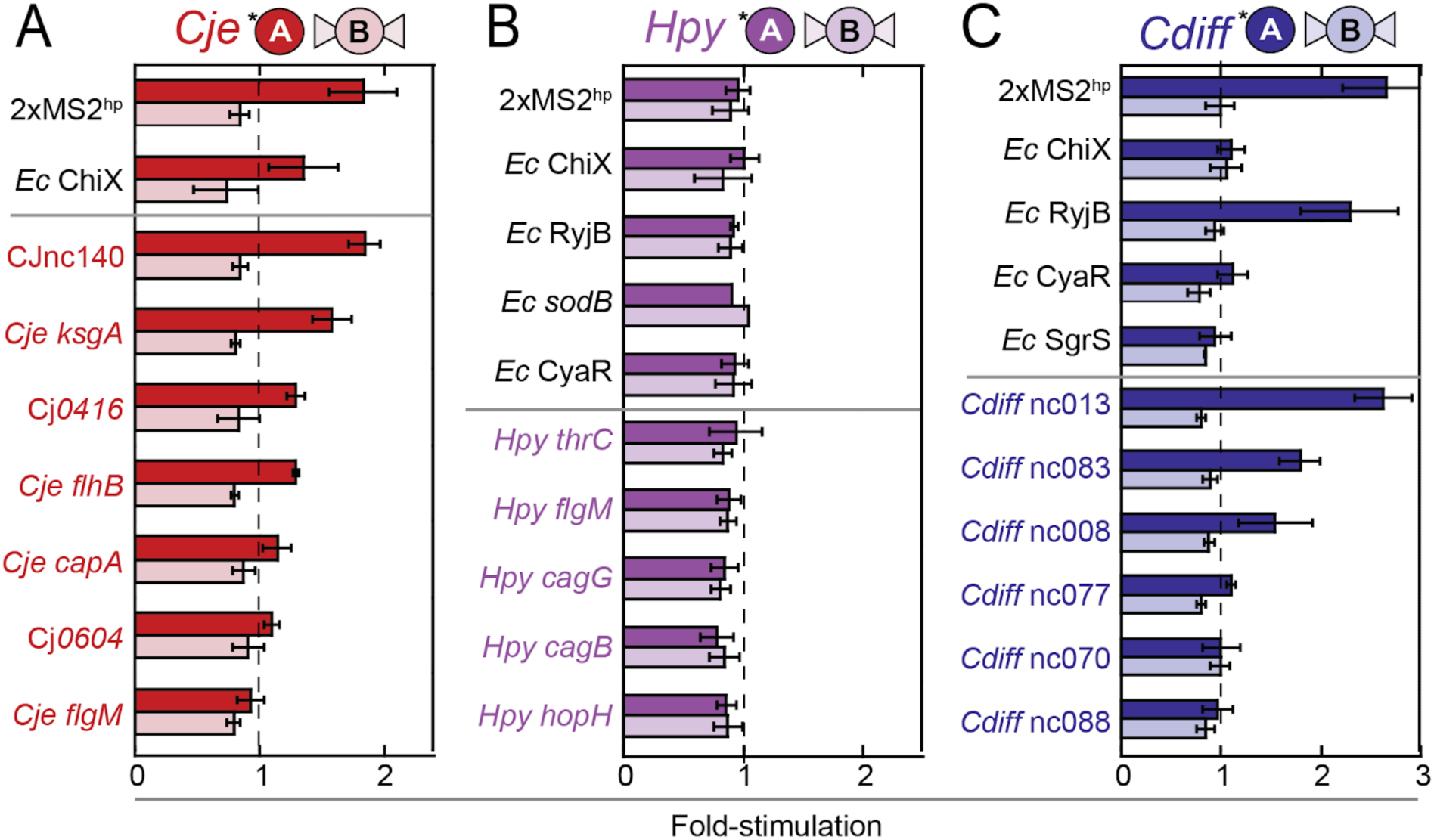
B3H detection of KhpA- and KhpB-RNA interactions with species-specific RNAs. Assays were performed in KB483 cells using pPrey plasmids containing KhpA (pKN29, pKN30, or pKN14) or KhpB (pCG112, pCG110, or pKN8) from each species or an α only negative control; pAdapter (pCW17); and the pBait-RNA panels described below. (A) *C. jejuni* Khp-RNA interactions. With the exception of the 2xMS2^hp^-only construct (pKB845), each hybrid RNA contained one MS2^hp^ followed by the indicated sRNA or mRNA fragment (pCH6, pNWL1, pNWL5, pNWL10, pNWL13, pNWL15, pNWL17, pNWL20) or corresponding negative controls expressing the MS2^hp^ moiety alone (pCH1, pSS1, pSS2). Certain RNAs were cloned into constructs containing a 5-bp GC clamp flanking the MS2^hp^ moiety (pSS1) or the MS2^hp^ moiety followed by a 7-bp GC clamp and *T_trpA_* terminator (pSS2); see Supplementary Figure S9 for details regarding matching control constructs. (B) *H. pylori* Khp–RNA interactions. Assays used *Hpy* KhpA or KhpB as prey proteins and bait RNAs containing *Hpy* RNAs (pNWL2, pNWL9, pNWL12, pNWL14, pNWL18), *E. coli* RNAs (pCH6, pSP25, pHL18, pLN35), or corresponding empty constructs. (C) *C. difficile* Khp–RNA interactions. Assays used *Cdiff* KhpA or KhpB as prey proteins and bait RNAs containing *Cdiff* RNAs (pKN21, pKN23–27), *E. coli* RNAs (pCH6, pSP25, pLN35, pLN91), or corresponding empty constructs.

Because these B3H data were collected with KhpA constructs possessing an extended flexible linker, we sought to determine if this modification influenced the observed RNA interactions, as it did for protein-protein interactions in the B2H assay (Supplementary Figure S2). We conducted side-by-side comparisons of pPrey-KhpA constructs with the original AAA linker or the extended (GGGGS)_3_-AAA linker. Even with the extended linker, *Hpy* KhpA failed to show any RNA interactions (Supplementary Figure S10), and very little difference was observed in the RNA interactions of *Cje* or *Cdiff* KhpA. This lack of impact in the B3H assay is consistent with the earlier finding that the strongest effects of the extended linker were mediated through the pBait(λCI) constructs, which are used only in the B2H assay.

### Direct comparison of Khp-RNA interactions across species

Given that KhpA from *C. difficile* and *C. jejuni* showed stronger interactions with RNAs selected from their respective species than *Hpy* KhpA did with its own RNAs, we asked whether this reflected intrinsic differences in the KhpA proteins themselves or differences in the RNA baits selected from the three species. We therefore compared the B3H interaction of each protein against the same panel of RNA baits, representing sRNAs and mRNA fragments from *E. coli, C. jejuni, H. pylori and C. difficile* that exhibited the strongest interactions in the species-specific panels above. This combined panel contained five sRNAs and three 5’UTRs/ORF fragments, with lengths ranging from 75-130 nucleotides and representing varying degrees of secondary structure (Supplementary Figure S11).

Results from these B3H assays largely mirrored the above results with species-specific RNAs: strong interactions were observed for KhpA proteins from *C. jejuni* (7/9 RNAs) and *C. difficile* (4/9 RNAs), but not for *Hpy* KhpA (0/9 RNAs) (Figure 6). While the *Cje* and *Cdiff* proteins each displayed some non-specific binding (interactions with RNAs from other species) the two proteins showed differences in the subset of RNAs with which they interact, suggesting some degree of specificity. Certain RNAs interacted with both *Cje* and *Cdiff* KhpA (the *Cdiff* RNA nc013 and the 2XMS2^hp^) whereas others only interacted with *Cje* KhpA (all four *Cje* and *Hpy* RNAs and *Ec* ChiX), and others only interacted with *Cdiff* KhpA (Cdiff nc083 and *Ec* RyjB). In contrast to the broad RNA interactions displayed by *Cdiff* KhpA and especially *Cje* KhpA, no B3H interactions were detected with the KhpB protein from any species.

**Figure 6.**
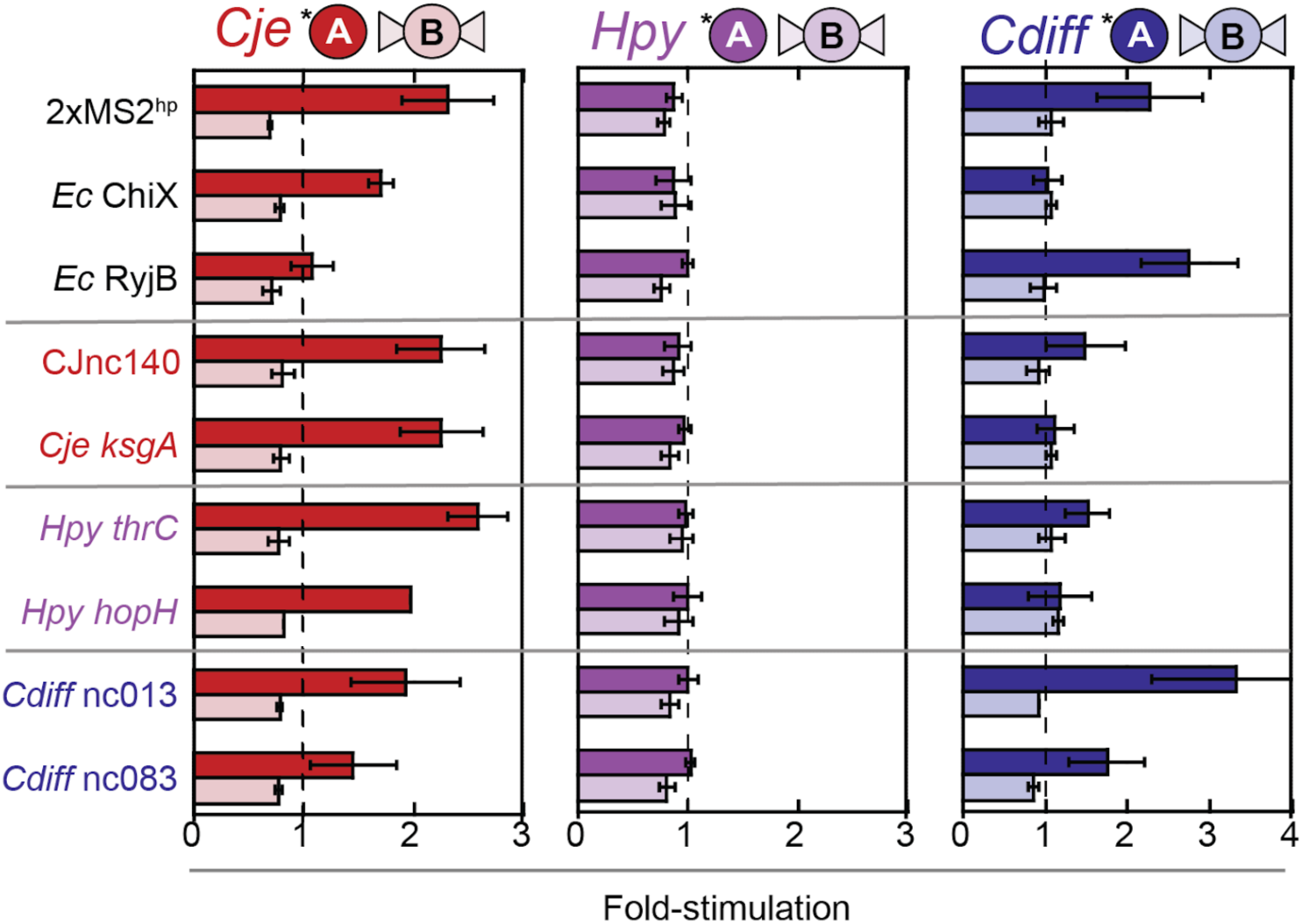
Comparative B3H analysis of KhpA- and KhpB-RNA interactions across species. B3H assays comparing RNA binding of KhpA and KhpB proteins from *C. jejuni* (red; pKN29, pCG112), *H. pylori* (purple; pKN30, pCG110) and *C. difficile* (blue; pKN14, pKN8) were performed as in Figure 5, but using a shared panel of RNAs representing strong interactors identified in the species-specific assays. Corresponding negative controls expressed the following: only the MS2^hp^ moiety (pCH1) for the 2xMS2^hp^–only construct (pKB845), *E. coli ChiX* (pCH6) and *RyjB* (pSP25); a 5-bp GC clamp flanking the MS2^hp^ moiety (pSS1) for CJnc140 (pNWL1) and *Cdiff* ncRNAs (pKN23, pKN27); and the MS2^hp^ moiety followed by a 7-bp GC clamp and T*_trpA_* terminator (pSS2) for *Cje* and *Hpy* mRNAs (pNWL2, pNWL12, pNWL17).

Given that isolated KH domains of KhpB demonstrated more robust protein-protein interactions than FL KhpB in the B2H assay (Supplementary Figure S4), we examined whether these isolated KH domains would exhibit better RNA interactions in the B3H system. We tested pPrey constructs containing the isolated KH domain of *Cje* and *Hpy* KhpB proteins against the same panel of 9 RNAs (as noted above, a functional analogous construct for the *Cdiff* protein could not be generated). Despite the strengthening of protein-protein interactions seen with the isolated KhpB KH domains, neither of these constructs displayed detectable RNA interaction with any of the 9 RNAs in the panel (Supplementary Figure S12). Thus, although the pPrey fusion proteins with full-length KhpB and the isolated KH domains showed robust heterodimerization in B2H assays with KhpA, no detectable protein-RNA interactions were observed for any construct of KhpB from the three species. Although KhpB from these species does not interact detectably with RNA on its own, it may nevertheless modulate the strength or specificity of KhpA interactions when partnering in a heterodimer.

### The GXXG motif is required for RNA binding by *Cje* and *Cdiff* KhpA

Given that the three KhpA orthologs differed in their RNA binding strength and specificity, we wanted to understand which structural features were responsible for these differences. Because residues within the GXXG motif (Figure 1A and Supplementary Figure S13) have been previously implicated in RNA binding (12, 42), we first examined the importance of this region in both *Cje* and *Cdiff* KhpA. We began by introducing two variants of the GXXG motif of *Cje* KhpA that have been utilized in prior studies: substitution of the inner residues with aspartate (GDDG) or of the outer residues with alanine (AXXA) (16, 42; Narayan et al., under review). Both substitutions in Cje KhpA (GKNG → GDDG or AKNA) eliminated detectable interaction with both the CJnc140 and Cdiff nc013 RNAs (Figure 7A, left).

**Figure 7.**
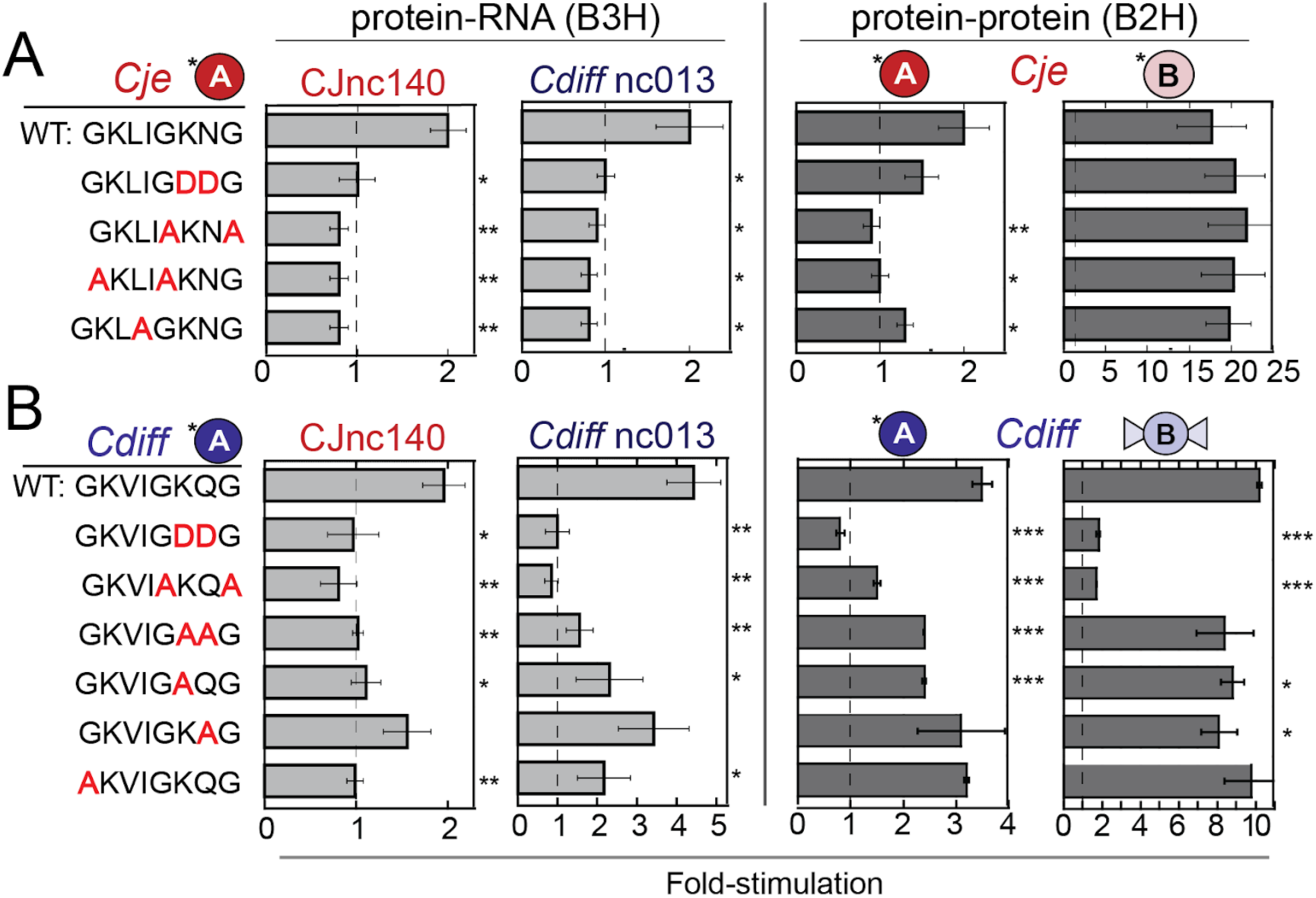
GXXG residues are required for KhpA-RNA interactions. In both panels, B3H assays (left) were performed in KB483 reporter cells to measure RNA binding of KhpA pPrey constructs to pBait–RNA constructs (pNWL1, pKN23) or a 1xMS2^hp^ negative-control construct (pCH1), while B2H assays (right) were performed in KB473 cells to measure protein dimerization of the same KhpA prey panel with pBait–protein constructs. (A) Analysis of *C. jejuni* GXXG motif variants. Assays utilized WT (pKN29) or mutant (pNWL21, pNWL22, pNWL28, pNWL29) *Cje* KhpA prey and the B3H pBait–RNA constructs listed above or B2H pBait constructs expressing *Cje* KhpA (pKN16) or the KhpB KH domain (pKB1293). (B) Analysis of *C. difficile* GXXG motif variants. Assays utilized WT (pKN14) or mutant (pKN32–37) *Cdiff* KhpA prey and the B3H pBait-RNA constructs listed above or B2H pBait constructs expressing *Cdiff* KhpA (pKN16) or full-length KhpB (pKN8). In both panels, asterisks indicate statistical significance of mutant interactions relative to WT (\**P* < 0.05, \*\**P* < 0.005, \*\*\**P* < 0.0005), as determined by Student’s t-tests.

Each *Cje* KhpA variant retained the ability to heterodimerize with the KH domain of KhpB in a B2H assay with WT-levels of efficiency (Figure 7A, right), indicating that the variants are still stably expressed and folded inside the cell. In contrast to this robust heterodimerization, KhpA homodimerization was impaired by each substitution, though only the AKNA variant produced a statistically significant decrease (see Discussion).

In addition to the “core” GXXG residues which are fully solvent exposed, most KhpA proteins possess an extended conserved structure: GXXIGXXG (Supplementary Figure S13). We were therefore curious whether upstream residues within this extended motif also play a role in RNA binding. The first of these variable residues is predicted to be solvent exposed, while the next one and isoleucine are predicted to face the core of the KH domain (Supplementary Figure S13B). We mutated the two upstream glycines in Cje KhpA to alanines (GKLIG → AKLIA) and found that this variant behaved similarly to the downstream GXXG substitutions: it exhibited a complete loss of RNA binding and KhpA homodimerization while retaining WT-levels of heterodimerization with the KhpB KH domain. Substitution of the conserved isoleucine to alanine (GKLIG → GKLAG) produced similar effects, confirming that residues across the extended motif of Cje KhpA play important roles in RNA binding and KhpA self-interaction, but not in interaction with KhpB.

To determine if the GXXG residues behave similarly in the *Cdiff* KhpA protein, we constructed an analogous set of point mutations. As before, substitution of the downstream GXXG residues (GDDG and AKQA) eliminated RNA binding and KhpA homodimerization (Figure 7B, left). However, unlike in the *Cje* protein, these *Cdiff* KhpA variants were no longer able to interact with the KhpB KH domain (Figure 7B, right). Without this control for the stability and folding of the variant proteins, the loss of RNA binding could not be definitively interpreted. To mitigate this, we made more conservative substitutions, replacing the two internal GXXG residues with alanines rather than aspartates (GAAG). This variant still lost RNA binding but was able to heterodimerize with the KhpB KH domain (Figure 7B). We also compared the relative contributions of the two internal residues by mutating them one at a time (GAQG and GKAG). B3H results showed that both residues contribute to RNA binding, with substitution of the lysine being more deleterious to RNA binding than substitution of the glutamine. Even substitution of a single upstream glycine to alanine was enough to impair RNA binding, emphasizing the importance of the overall conformation of this extended GXXG motif within KhpA. Together, the above results demonstrate that both the “upstream” and “downstream” residues in the extended GXXIGXXG motif contribute to the RNA-binding activity of *Cje* and *Cdiff* KhpA.

### The GXXG motif of *Hpy* KhpA contributes to its weak interaction with RNA

Given that *Hpy* KhpA did not interact with any RNA tested, we asked whether the sequence of its GXXG motif accounted for its lack of binding. Of the three KhpA orthologs investigated here, each possesses a positively charged lysine in the first variable position of the GXXG motif, but they differ in the second variable residue: the two KhpA proteins that displayed RNA interactions in the B3H assay (*Cje* and *Cdiff*) both contain a neutral polar residue in the second position (KN or KQ), while *Hpy* KhpA — which did not demonstrate RNA binding — contains a negatively charged residue in this position (KE). We hypothesized that this difference in sequence and electrostatics of the GXXG motif might account for why the *Cje* and *Cdiff* orthologs were able to bind RNA in the absence of their KhpB partners, while *Hpy* KhpA was not.

To test this, we asked whether substituting the GXXG residues in *Cje* and *Cdiff* KhpA with the corresponding *Hpy* residues would be sufficient to abolish RNA binding. Substitution of the downstream GXXG residues in Cje KhpA with the corresponding Hpy residues (GKNG → GKEG) led to a partial — though not statistically significant — reduction in RNA interaction and homodimerization, without altering heterodimerization efficiency (Figure 8A). Substitution of the upstream GXXG residues with Hpy residues (GKLIG → GHVIG) fully eliminated RNA binding to all three RNAs tested (CJnc140, Hpy *thrC* and Cdiff nc013) as well as homodimerization, without altering interaction with the KhpB KH domain, as did simultaneous substitution of both upstream and downstream residues. In contrast, replacing the Cje KhpA residues with those from the RNA-binding Cdiff ortholog (GKNG → GKQG) preserved full interaction with all three RNAs, as well as with KhpA and KhpB.

**Figure 8.**
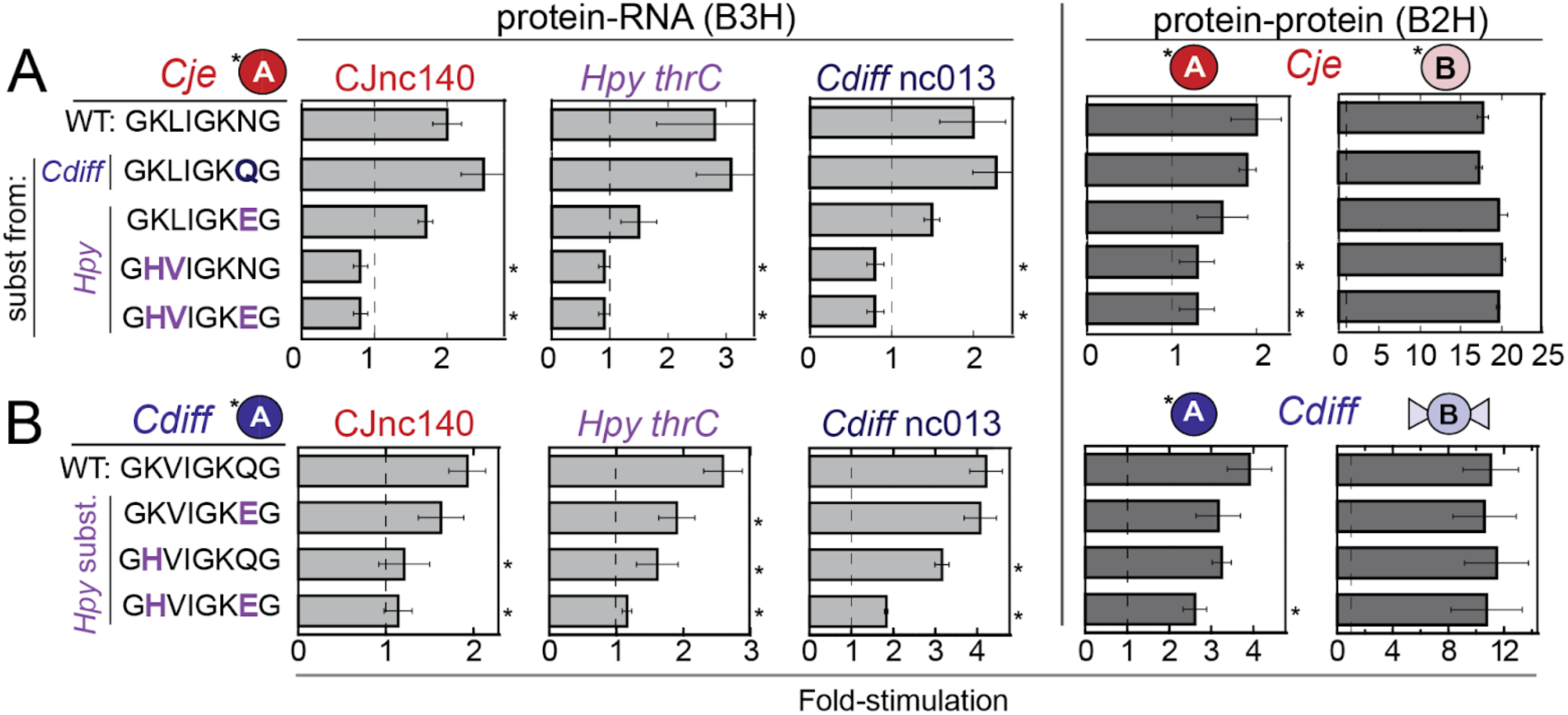
Introduction of *Hpy* GXXG residues is sufficient to disrupt *Cje* and *Cdiff* KhpA-RNA interactions. In both panels, B3H assays (left) were performed in KB483 reporter cells to measure RNA binding of KhpA pPrey constructs to pBait–RNA constructs (pNWL1, pNWL2, pKN23) or a 1xMS2^hp^ negative-control construct (pCH1), while B2H assays (right) were performed in KB473 cells to measure protein dimerization of the same KhpA prey panel with pBait–protein constructs. (A) Analysis of *Cje* KhpA variants containing *Hpy*- and *Cdiff*-type GXXG residues. Assays utilized WT (pKN29) or mutant (pNWL23, pNWL24, pNWL30, pNWL31) *Cje* KhpA prey and the B3H pBait–RNAs listed above or B2H pBait constructs expressing *Cje* KhpA (pKN16) or the KhpB KH domain (pKB1293). (B) Analysis of *Cdiff* KhpA substituted with *Hpy*-type GXXG residues. Assays utilized WT (pKN14) or mutant (pNWL34–36) *Cdiff* KhpA prey and the B3H pBait–RNAs listed above or B2H pBait constructs expressing *Cdiff* KhpA (pKN16) or full-length KhpB (pKN8). In both panels, asterisks indicate statistical significance of mutant interactions relative to WT (\**P* < 0.05, \*\**P* < 0.005, \*\*\**P* < 0.0005), as determined by Student’s t-tests.

Similar effects were observed when *Cdiff* KhpA residues were substituted with the corresponding *Hpy* residues: substitution of the downstream GXXG residues alone resulted in a partial reduction of interaction with *Hpy thrC*, while substitution of the upstream GXXG residues had stronger effects against all three RNAs tested (Figure 8B). Interestingly, an additive effect of substituting the upstream and downstream *Hpy* residues was observed in the interaction of these variants with *Cdiff* nc013. Taken together, these results indicate that the difference in the GXXG residues of the three KhpA proteins indeed accounts for the lack of RNA interaction by *Hpy* KhpA.

Given that the *Hpy* GXXG residues were sufficient to disrupt RNA binding by *Cdiff* or *Cje* KhpA, we next explored whether the introduction of *Cdiff* or *Cje* KhpA GXXG residues would be sufficient to restore RNA-binding activity to *Hpy* KhpA. While substitution of the upstream and/or downstream GXXG residues from either species did not interfere with *Hpy* KhpA’s ability to interact with itself or with KhpB, none of the variants produced detectable RNA interaction (Supplementary Figure S14). This indicates that *Hpy* KhpA likely lacks at least one additional RNA-binding determinant beyond its GXXG residues.

## DISCUSSION

This study provides the first systematic comparison of the KhpA and KhpB proteins across multiple bacterial species. By heterologously expressing these proteins in *E. coli* reporter cells, we were able to compare the molecular interactions of *C. jejuni*, *H. pylori*, and *C. difficile* orthologs side by side. Across all species, homodimerization was more prevalent for KhpA than for KhpB, and KhpA/B heterodimerization was favored over homodimerization of either protein. While KhpA proteins from *C. jejuni* and *C. difficile* showed robust RNA-binding activity, the *H. pylori* KhpA and all three of the KhpB proteins did not interact with any of the RNAs in our assays. We further show that the GXXG motifs of KhpA are critical for both RNA interaction and homodimerization. Taken together, these results suggest that while the heterodimeric architecture of the KhpA/B complex is evolutionarily conserved, the molecular determinants of RNA recognition have diverged between bacterial species.

### Molecular determinants of Khp protein-protein interactions

Consistent with previous studies in other species (13, 16, 19, 21; Narayan et al., under review), we found that KhpA and KhpB form strong heterodimers across the three species investigated here. Our detection of KhpA homodimers for all three orthologs is also consistent with previous findings in *S. pneumoniae* (13). While a recent crystallization study using *T. thermophilus* KhpB suggests this protein has the potential to homodimerize (14), our B2H experiments did not detect any homodimerization of full-length KhpB proteins. However, we found that the isolated KH domains of *Cje* and *Hpy* KhpB were able to homodimerize to a similar extent as KhpA proteins (Supplementary Figure S4). These results suggest that the additional Jag-N and/or R3H domains of KhpB may inhibit the protein-protein interactions of its KH domain and could play a role in tuning the equilibrium of dimeric species. However, it remains to be seen if these inhibitory effects are influenced by geometric constraints of the B2H fusion proteins or if they occur with native, unfused proteins.

Our findings also demonstrate that while the location of the heterodimerization interface of KhpA and KhpB is structurally conserved across species, the specific residues driving interaction specificity have evolved differently. We observed a notable lack of reciprocity in cross-species interactions: *Cje* KhpA interacted strongly with *Hpy* KhpB, but *Cje* KhpB did not interact with *Hpy* KhpA (Supplementary Figure 3). This asymmetry can be rationalized through structural predictions of the interface: lysine residues positioned across from one another in the *Hpy* KhpA/*Cje* KhpB pairing would create steric and electrostatic clash, whereas smaller aspartate and glutamate residues in the *Cje* KhpA/*Hpy* KhpB pairing would be better accommodated. These cross-species heterodimer interactions demonstrate how the KH-domain interface can be tuned through small changes in the amino acids at the dimer interface.

### RNA interactions of KhpA proteins

Because our *E. coli* reporter strain lacks native Khp orthologs, any RNA interactions detected in the B3H assay must arise from KhpA or KhpB acting independently — either as monomers or homodimers. This allowed us to investigate the independent, intrinsic RNA-binding capacity of each protein. Our comparative B3H data reveal that *Cje* and *Cdiff* KhpA orthologs can bind RNA independently, whereas *Hpy* KhpA did not interact with any of the tested RNAs.

While both *Cje* and *Cdiff* KhpA proteins bound RNA, they showed distinct RNA ligand preferences: *Cje* KhpA bound broadly to sRNAs and mRNA fragments from all species investigated while *Cdiff* KhpA showed a narrower preference for sRNAs from its own species or from *E. coli* (Figures 5 and 6). Notably, we found no striking pattern between the sequences or predicted secondary structures of the RNA baits that interacted with each protein and those that did not (Supplementary Figures S5-S8). This lack of an obvious motif, combined with the observation that neither *Cje* nor *Cdiff* KhpA showed a strict preference for RNAs from their own species, supports a view of the Khp proteins as global RNA binding proteins, as suggested by prior transcriptome-wide analyses across numerous species (15–20)

Our B3H results for *C. jejuni* KhpA generally align with our unpublished findings from global RIP-seq analyses conducted in its native species. Specifically, of the four *Cje* RNAs for which we detected B3H interactions, three — *ksgA*, *Cj0416*, and *flhB* — were previously detected to co-immunoprecipitate with KhpA even in the absence of KhpB (Narayan et al., under review). The fourth, CJnc140, was found to be just below the threshold of statistical significance for enrichment in the KhpA-IP. Together, these B3H and RIP-seq data suggest that KhpA monomers or homodimers can mediate these RNA interactions independent of KhpB, at least for some RNAs.

The relationship between our B3H data and prior *in vivo* studies is more challenging to interpret for the *C. difficile* proteins, as the native RIP-seq and co-IP studies were conducted in a wild-type background where both KhpA and KhpB were present (Lamm-Schmidt et al. 2021).

Of the three RNAs that co-immunoprecipitated with *Cdiff* KhpA in its native species (nc008, nc070, and nc088), our B3H assay detected an interaction only with nc008. This suggests that while nc008 can be recognized by KhpA alone, other native ligands may require a KhpA/B heterodimer for stable binding in the cell.

### Potential coupling of KhpA homodimerization and RNA binding

An interesting question from our B3H data is whether the observed RNA interactions arise from a monomeric or homodimeric form of KhpA. While this is challenging to determine definitively, our mutagenesis results suggest that KhpA may interact with RNA as a homodimer in the B3H system. For GXXG motif substitutions in both *Cje* and *Cdiff* KhpA proteins, the reduction of KhpA-KhpA homodimer interactions (B2H) largely mirrors the loss of KhpA-RNA interactions (B3H) relative to the WT protein (Figures 7 and 8). Notably, this trend is much more pronounced for KhpA homodimerization than for its interaction with KhpB, which remained stable across most variants. The heterodimer interactions serve as an important control, indicating that most substitutions in KhpA specifically disrupted homodimerization and RNA binding rather than causing general instability or misfolding of the protein.

The mirrored relationship between KhpA homodimerization and RNA interaction strength across mutants suggests that these activities are strongly coupled inside the reporter cell. That is, KhpA’s interaction with RNA may be stabilized by homodimerization and, conversely, homodimerization may be stabilized by binding to cellular RNA. Notably, the architecture of the B3H assay presents two copies of the α-NTD-prey proteins per RNAP molecule (Figure 4A) potentially facilitating KhpA homodimerization within the complex.

### The GXXG motif as a key determinant of RNA binding

Our site-directed mutagenesis results demonstrate the importance of the GXXG motif for RNA binding by KhpA proteins. Consistent with the deleterious *in vivo* effects of GDDG and/or AXXA substitutions in *S. pneumoniae* and *C. jejuni* (16; Narayan et al., under review), both of these substitutions were deleterious for the RNA-binding activity of *Cje* and *Cdiff* KhpA in B3H experiments. In addition, our mutagenesis data extend beyond previous studies in several ways. First, we showed that replacing the internal GXXG residues with alanine (GAAG) is sufficient to disrupt RNA interactions. This suggests that the loss of activity previously observed in GDDG mutants is not simply a dominant-negative effect from electrostatic repulsion of the aspartate residues, but a consequence of losing specific side chains required for the interaction. Second, by isolating the individual effects of the two variable positions in *Cdiff* KhpA’s GXXG motif (GKQG), we demonstrated that the lysine contributes more to RNA interaction than the glutamine, consistent with the lysine’s positive charge. Furthermore, we showed that the conserved glycine and isoleucine residues upstream of the core GXXG motif are both important for KhpA-RNA interactions. Given that this isoleucine is predicted to point back into the core of the KhpA monomer rather than being solvent exposed (Supplementary Figure S13), its contribution to RNA binding suggests that the overall conformation of this region is critical to positioning residues that directly interact with RNA.

Our findings also suggest the lack of RNA interaction by the *Hpy* KhpA ortholog is driven by its specific GXXG sequence, which is less electropositive than that of *Cje* or *Cdiff* KhpA. While the GXXG motif residues of *Cje* or *Cdiff* KhpA were not sufficient to confer RNA binding activity to the *Hpy* KhpA protein (Supplementary Figure S14), inserting the GXXG residues of *Hpy* KhpA into the *Cje* and *Cdiff* proteins completely eliminated their ability to bind RNA (Figure 8). Together, these results provide strong evidence that the GXXG motif, while not the sole determinant, plays a critical role in shaping RNA-binding activity,

### Divergent roles of KhpA and KhpB in RNA binding

The three KhpB orthologs investigated here differ in their conservation of canonical sequence elements: the *Cdiff* protein has intact R3H and GXXG motifs, while both the *Cje* and *Hpy* KhpB proteins lack an R3H motif and the *Cje* GXXG motif is also degenerate, containing an alanine at one of the two glycine positions (Figure 1A). Despite these differences, our B3H experiments produced the same results for all three proteins: no RNA interactions were detectable for either the FL or isolated KH domains, even while these same fusion proteins showed robust B2H interactions. This stands in contrast to prior findings suggesting a role for KhpB in RNA binding in certain species. In *F. nucleatum*, KhpB appears to play a dominant role in RNA binding — interacting with RNA *in vitro* even when KhpA did not — though its affinity increased in the presence of KhpA (20). In addition, *T. thermophilus* KhpB was recently found to bind RNA on its own with micromolar affinity (14), but the corresponding KhpA protein was not able to be expressed for comparison. While *Cdiff* KhpB was previously found to bind widely to cellular RNA *in vivo* (15), these data were collected in a wild-type background where KhpA was present, so some of the interactions may depend on the presence of KhpA within a heterodimer. In alignment with our B3H findings that KhpA appears to be the primary driver of independent RNA interactions for the *Cje* and *Cdiff* proteins, a recent study showed that *D. radiodurans* KhpA binds RNA *in vitro* with higher affinity than KhpB (21).

### Methodological considerations and updated B2H/B3H tools

This study provides updated tools and new precedents for the B2H and B3H assays. The introduction of extended flexible linkers into pBait and pPrey constructs proved helpful in detecting protein-protein interactions. Such linkers may be especially useful for proteins where the N-terminus is involved in the binding interface, providing an alternative to C-terminal fusions to the ω subunit of RNAP (43, 44). Interestingly, the restrictions of the original triple-alanine linker seemed to impact the λCI fusions (pBait) more significantly than the α-NTD fusions (pPrey). Consistent with this, we observed large effects of the extended linker on B2H (protein-protein) interactions but minimal changes in B3H (RNA-protein) interactions, which utilize an RNA bait molecule rather than a bait protein directly fused to λCI.

This work shows that the *E. coli*-based B3H assay provides a powerful platform for investigating interactions of diverse bacterial orthologs, away from the influence of species-specific endogenous RNAs and interacting/competing proteins. While previous studies encountered challenges purifying soluble Khp proteins (14, 20), the fusion moieties in our system may stabilize these proteins, allowing for a more complete comparison of their RNA-binding activities. In addition, the dual nature of the B2H/B3H assays — utilizing identical reporter strains and prey fusion proteins — provides a built-in validation system: the fact that all pPrey-KhpA and -KhpB fusion proteins exhibited strong interactions in B2H assays serves as an important control for proper protein folding and expression. This validation strengthens the interpretation of negative B3H results suggesting that *Hpy* KhpA and the three KhpB orthologs likely lack high-affinity RNA binding on their own. At the same time, the B3H assay’s detection limit (45) may preclude the observation of weaker interactions such as the micromolar binding affinities reported for isolated KhpB proteins *in vitro* (14, 20, 21).

### Outlook

Together, the results of this study and prior investigations reveal a complex landscape where the relative contributions of KhpA and KhpB to RNA binding vary substantially across species. While we found that KhpA/B heterodimerization and KhpA homodimerization are relatively consistent features across orthologs, the RNA-binding capacity of KhpA proteins varies considerably — even between the closely related ε-proteobacteria *C. jejuni* and *H. pylori*. This supports a model of a dynamic evolutionary process that tunes RNA-binding activity within each organism. While this study focuses on isolated KhpA and KhpB subunits, it is possible that more similarities will emerge when RNA binding of the heterodimeric complex is interrogated. Given that eukaryotic KH-domain proteins often utilize multivalent interactions of multiple KH domains within a single polypeptide (26), it is striking that KhpA/B heterodimers are so dominant both in this and prior studies.

A critical next step for the B3H assay is therefore to develop expression vectors for the simultaneous co-expression of two heterologous proteins in order to examine the effect of an untagged protein on the RNA-binding behavior of a prey fusion protein. We hypothesize that the addition of KhpB to KhpA binding experiments may tune the strength and specificity of *Cje* and *Cdiff* KhpA proteins and that the presence of KhpA may be required for interactions of certain KhpB proteins to be detected. It will be interesting in the future to determine the role that different KhpB domains play in RNA-binding in the context of the KhpA/B heterodimers.

In conclusion, this work emphasizes that as Khp proteins continue to be studied across a range of bacterial species, it is important to avoid the assumption that observations in one species will necessarily apply to another. The modularity of the domains within KhpA and KhpB allows for the evolution of distinct behaviors such as tuning the equilibrium between monomers and dimers, recruiting additional factors, or adjusting the strength and specificity of RNA interactions. Each of these features is likely to play an important role in the mechanisms by which KhpA and KhpB regulate bacterial post-transcriptional gene expression.

## SUPPLEMENTARY MATERIAL

Supplementary materials for this article are available online, including Supplementary Tables S1 to S3 and Supplementary Figures S1 to S14.

## DATA AVAILABILITY STATEMENT

The data underlying this article are available in the article and in its online supplementary materials.

## ACKNOWLEDGEMENTS

We thank members of the Berry Lab and Amy Camp for helpful discussions and feedback and Annie Williams for assistance with cloning as well as Philipp Kible for isolation of bacterial genomic DNA and Sara Eisenbart for help with RIP-seq experiments.

## FUNDING

This work was supported by the National Institutes of Health [2R15GM135878 to K.E.B.], the Camille and Henry Dreyfus Foundation, and Mount Holyoke College.

## AUTHOR CONTRIBUTIONS

K.T.N., N.W.L. C.M.G., S.J. and Y.S. designed and cloned plasmids and tested them in B3H assays. K.E.B. conceived of the project, obtained funding, and oversaw the experimental implementation. M.N., S.S. and C.M.S. provided genomic DNA and RIP-seq candidates. K.T.N, N.W.L. and K.E.B. analyzed data and completed statistical analyses. K.T.N. and K.E.B made the figures and wrote the manuscript with input and feedback from N.W.L, C.M.G., S.J., Y.S., M.N. and C.M.S.

## Notes

### Competing Interest Statement

The authors have declared no competing interest.

## REFERENCES

1. Papenfort, K. and Melamed, S. (2023) Small RNAs, Large Networks: Posttranscriptional Regulons in Gram-Negative Bacteria. Annu. Rev. Microbiol., 77, 23–43.

2. Wagner, E.G.H. and Romby, P. (2015) Small RNAs in bacteria and archaea: who they are, what they do, and how they do it. Adv. Genet., 90, 133–208.

3. Svensson, S.L. and Sharma, C.M. (2016) Small RNAs in Bacterial Virulence and Communication. Microbiol. Spectr., 4, 4.3.02.

4. Ng Kwan Lim, E., Sasseville, C., Carrier, M.-C. and Massé, E. (2021) Keeping up with RNA-based regulation in bacteria: New roles for RNA binding proteins. Trends Genet., 37, 86–97.

5. Bobrovskyy, M., Vanderpool, C.K. and Richards, G.R. (2015) Small RNAs Regulate Primary and Secondary Metabolism in Gram-negative Bacteria. Microbiol. Spectr., 3.

6. Updegrove, T.B., Zhang, A. and Storz, G. (2016) Hfq: the flexible RNA matchmaker. Curr. Opin. Microbiol., 30, 133–138.

7. Olejniczak, M. and Storz, G. (2017) ProQ/FinO-domain proteins: another ubiquitous family of RNA matchmakers? Mol. Microbiol., 104, 905–915.

8. Holmqvist, E., Berggren, S. and Rizvanovic, A. (2020) RNA-binding activity and regulatory functions of the emerging sRNA-binding protein ProQ. Biochim. Biophys. Acta BBA - Gene Regul. Mech., 1863, 194596.

9. Quendera, A.P., Seixas, A.F., dos Santos, R.F., Santos, I., Silva, J.P.N., Arraiano, C.M. and Andrade, J.M. (2020) RNA-binding proteins driving the regulatory activity of small non-coding RNAs in bacteria. Front. Mol. Biosci., 7, 78.

10. Olejniczak, M., Jiang, X., Basczok, M.M. and Storz, G. (2021) KH domain proteins: Another family of bacterial RNA matchmakers? Mol. Microbiol., 00, 1–10.

11. Grishin, N. (1998) The R3H motif: a domain that binds single-stranded nucleic acids. Trends Biochem. Sci., 23, 329–330.

12. Valverde, R., Edwards, L. and Regan, L. (2008) Structure and function of KH domains: Structure and function of KH domains. FEBS J., 275, 2712–2726.

13. Winther, A.R., Kjos, M., Stamsås, G.A., Håvarstein, L.S. and Straume, D. (2019) Prevention of EloR/KhpA heterodimerization by introduction of site-specific amino acid substitutions renders the essential elongasome protein PBP2b redundant in *Streptococcus pneumoniae*. Sci. Rep., 9, 3681.

14. Fukui, K., Murakawa, T., Baba, S., Kumasaka, T. and Yano, T. (2025) KH–R3H domain cooperation in RNA recognition by the global RNA-binding protein KhpB. Nat. Commun., 16, 8028.

15. Lamm-Schmidt, V., Fuchs, M., Sulzer, J., Gerovac, M., Hör, J., Dersch, P., Vogel, J. and Faber, F. (2021) Grad-seq identifies KhpB as a global RNA-binding protein in *Clostridioides difficile* that regulates toxin production. microLife, 2, uqab004.

16. Zheng, J.J., Perez, A.J., Tsui, H.-C.T., Massidda, O. and Winkler, M.E. (2017) Absence of the KhpA and KhpB (JAG/EloR) RNA-binding proteins suppresses the requirement for PBP2b by overproduction of FtsA in *Streptococcus pneumoniae* D39: KhpA/B regulation of pneumococcal FtsA expression. Mol. Microbiol., 106, 793–814.

17. Hör, J., Garriss, G., Di Giorgio, S., Hack, L., Vanselow, J.T., Förstner, K.U., Schlosser, A., Henriques-Normark, B. and Vogel, J. (2020) Grad-seq in a Gram-positive bacterium reveals exonucleolytic SRNA activation in competence control. EMBO J., 39, e103852.

18. Riediger, M., Spät, P., Bilger, R., Voigt, K., Maček, B. and Hess, W.R. (2021) Analysis of a photosynthetic cyanobacterium rich in internal membrane systems via gradient profiling by sequencing (Grad-seq). Plant Cell, 33, 248–269.

19. Michaux, C., Gerovac, M., Hansen, E.E., Barquist, L. and Vogel, J. (2023) Grad-seq analysis of *Enterococcus faecalis* and *Enterococcus faecium* provides a global view of RNA and protein complexes in these two opportunistic pathogens. microLife, 4, uqac027.

20. Zhu, Y., Ponath, F., Cosi, V. and Vogel, J. (2024) A global survey of small RNA interactors identifies KhpA and KhpB as major RNA-binding proteins in *Fusobacterium nucleatum*. Nucleic Acids Res., 52, 3950–3970.

21. Han, R., Quinones-Diaz, B.I., Cordova, A., Niese, B., Sweet, P., Fang, J., Engels, S., Chen, A., Gordon, V.D. and Contreras, L.M. (2025) RNA Binding Proteins KhpA and KhpB Interact with Small Regulatory 2 RNAs and Affect Global Gene Expression in Deinococcus radiodurans. BioRxiv, 10.1101/2025.09.18.676964.

22. Stamsås, G.A., Straume, D., Ruud Winther, A., Kjos, M., Frantzen, C.A. and Håvarstein, L.S. (2017) Identification of EloR (Spr1851) as a regulator of cell elongation in *Streptococcus pneumoniae*: Regulation of cell elongation in *S. pneumoniae*. Mol. Microbiol., 105, 954–967.

23. Ulrych, A., Holečková, N., Goldová, J., Doubravová, L., Benada, O., Kofroňová, O., Halada, P. and Branny, P. (2016) Characterization of pneumococcal Ser/Thr protein phosphatase *phpP* mutant and identification of a novel PhpP substrate, putative RNA binding protein Jag. BMC Microbiol., 16, 247.

24. Myrbråten, I.S., Wiull, K., Salehian, Z., Håvarstein, L.S., Straume, D., Mathiesen, G. and Kjos, M. (2019) CRISPR interference for rapid knockdown of essential cell cycle genes in *Lactobacillus plantarum*. mSphere, 4, e00007-19.

25. Van Opijnen, T. and Camilli, A. (2012) A fine scale phenotype–genotype virulence map of a bacterial pathogen. Genome Res., 22, 2541–2551.

26. Nicastro, G., Taylor, I.A. and Ramos, A. (2015) KH–RNA interactions: back in the groove. Curr. Opin. Struct. Biol., 30, 63–70.

27. Winther, A.R., Kjos, M., Herigstad, M.L., Håvarstein, L.S. and Straume, D. (2021) EloR Interacts with the Lytic Transglycosylase MltG at Midcell in Streptococcus pneumoniae R6. J. Bacteriol., 203, e00691–20.

28. Burnham, P.M. and Hendrixson, D.R. (2018) Campylobacter jejuni: collective components promoting a successful enteric lifestyle. Nat. Rev. Microbiol., 16, 551–565.

29. Guery, B., Galperine, T. and Barbut, F. (2019) *Clostridioides difficile* : diagnosis and treatments. BMJ, 10.1136/bmj.l4609.

30. Salama, N.R., Hartung, M.L. and Müller, A. (2013) Life in the human stomach: persistence strategies of the bacterial pathogen Helicobacter pylori. Nat. Rev. Microbiol., 11, 385–399.

31. Pandey, S., Gravel, C.M., Stockert, O.M., Wang, C.D., Hegner, C.L., LeBlanc, H. and Berry, K.E. (2020) Genetic identification of the functional surface for RNA binding by *Escherichia coli* ProQ. Nucleic Acids Res., 48, 4507–4520.

32. Berry, K.E. and Hochschild, A. (2018) A bacterial three-hybrid assay detects *Escherichia coli* Hfq-sRNA interactions *in vivo*. Nucleic Acids Res., 46, e12.

33. Nguyen, L.D., LeBlanc, H. and Berry, K.E. (2025) Improved constructs for bait RNA display in a bacterial three-hybrid assay. Sci. Rep., 15, 3820.

34. Stockert, O.M., Gravel, C.M. and Berry, K.E. (2022) A bacterial three-hybrid assay for forward and reverse genetic analysis of RNA–protein interactions. Nat. Protoc., 17, 941–961.

35. Kerpedjiev, P., Hammer, S. and Hofacker, I.L. (2015) Forna (force-directed RNA): Simple and effective online RNA secondary structure diagrams. Bioinformatics, 31, 3377–3379.

36. Lorenz, R., Bernhart, S.H., Höner Zu Siederdissen, C., Tafer, H., Flamm, C., Stadler, P.F. and Hofacker, I.L. (2011) ViennaRNA Package 2.0. Algorithms Mol. Biol., 6, 26.

37. Abramson, J., Adler, J., Dunger, J., Evans, R., Green, T., Pritzel, A., Ronneberger, O., Willmore, L., Ballard, A.J., Bambrick, J., et al. (2024) Accurate structure prediction of biomolecular interactions with AlphaFold 3. Nature, 630, 493–500.

38. Dove, S.L., Joung, J.K. and Hochschild, A. (1997) Activation of prokaryotic transcription through arbitrary protein-protein contacts. Nature, 386, 627–630.

39. Dove, S.L. and Hochschild, A. (2004) A Bacterial Two-Hybrid System Based on Transcription Activation. In Protein-Protein Interactions. Humana Press, New Jersey, Vol. 261, pp. 231–246.

40. Stein, E.M., Kwiatkowska, J., Basczok, M.M., Gravel, C.M., Berry, K.E. and Olejniczak, M. (2020) Determinants of RNA recognition by the FinO domain of the *Escherichia coli* ProQ protein. Nucleic Acids Res., 10.1093/nar/gkaa497.

41. Dugar, G., Herbig, A., Förstner, K.U., Heidrich, N., Reinhardt, R., Nieselt, K. and Sharma, C.M. (2013) High-Resolution Transcriptome Maps Reveal Strain-Specific Regulatory Features of Multiple Campylobacter jejuni Isolates. PLoS Genet., 9, e1003495–15.

42. Hollingworth, D., Candel, A.M., Nicastro, G., Martin, S.R., Briata, P., Gherzi, R. and Ramos, A. (2012) KH domains with impaired nucleic acid binding as a tool for functional analysis. Nucleic Acids Res., 40, 6873–86.

43. Dove, S.L. and Hochschild, A. (1998) Conversion of the v subunit of Escherichia coli RNA polymerase into a transcriptional activator or an activation target. Genes Dev., 12, 745–754.

44. Vallet-Gely, I., Donovan, K.E., Fang, R., Joung, J.K. and Dove, S.L. (2005) Repression of phase-variable *cup* gene expression by H-NS-like proteins in *Pseudomonas aeruginosa*. Proc. Natl. Acad. Sci., 102, 11082–11087.

45. Wang, C.D., Mansky, R., Leblanc, H., Gravel, C.M. and Berry, K.E. (2021) Optimization of a bacterial three-hybrid assay through *in vivo* titration of an RNA–DNA adapter protein. RNA, 27, 513–526.

