## Supplementary Materials for "Molecular genetic characterization of bacterial KH-domain proteins"

This pdf file includes:

Supplementary Tables S1 to S3

Supplementary Figures S1 to S14

Supplementary References

**Supplementary Table S1. Bacterial strains used in this study.**

| **Strain** | **Genotype** | **Antibiotic Resistance** | **Source** |
|---|---|---|---|
| NEB5α-F′Iq | *E. coli lacIq host strain for plasmid construction.* | TetR | New England Biolabs |
| KB483 | FW102 Δhfq::kan harboring an F′ episome bearing test promoter plac-OL2-62 fused to lacZ. | KanR; TetR; StrR | (Pandey et al. 2020) |
| KB473 | FW102 Δhfq (unmarked) harboring an F' kan bearing test promoter placOL2-62 fused to lacZ. | KanR; TetR; StrR | (Berry and Hochschild 2018) |
| *Campylobacter jejuni* NCTC11168 WT | *C. jejuni* strain from which gDNA served as template for PCR of prey proteins and bait RNAs | | Sharma Lab |
| *Helicobacter pylori* G27 WT | *H. pylori* strain from which gDNA served as template for PCR of prey proteins and bait RNAs | | Sharma Lab |
| *Clostridioides difficile* strain 630 | *Clostridioides difficile s*train from which gDNA served as template for PCR of prey proteins and bait RNAs | | ATCC #BAA-1382D-5 |

**Supplementary Table S2. Plasmids used in this study.**

| Plasmid Name | Description | Details | Source or construction in this study |
|---|---|---|---|
| p35u4 (Addgene #174786) | pAdapter: pAC-$p_{constit}$-CI-MS2$^{CP}$ | Encodes CI-MS2CP fusion protein under control of a constitutive promoter; p15A origin of replication; confers CmR | (Wang et al. 2021) |
| pACλCI (Addgene #53730) | pACλCI empty vector | Encodes full-length λCI under the control of the lacUV5 promoter; confers CmR | (Dove et al. 1997) |
| pBr-α (Addgene #53731) | pBR-α-empty vector | Encodes residues 1-248 of the alpha vector fused under control of lpp and lacUV5 promoters; confers AmpR | (Dove et al. 1997) |
| pCG110 | pPrey-α-*Hpy*-KhpB | Encodes residues 1-248 of the alpha subunit of RNA polymerase fused via three alanine residues to full-length *Helicobacter pylori* KhpB; confers AmpR | Backbone: pKB817<br>insert: oSJ7 + oNWL13 |
| pCG111 | pBait-CI-*Hpy*-KhpB | Encodes residues 1-236 of λCI fused via three alanine residues to full-length *Helicobacter pylori* KhpB; confers CamR | Backbone: pKB816<br>insert: oSJ7 + oNWL13 |
| pCG112 | pPrey-α-*Cje*-KhpB | Encodes residues 1-248 of the alpha subunit of RNA polymerase fused via three alanine residues to full-length *Campylobacter jejuni* KhpB; confers AmpR | Backbone: pKB817<br>insert: oKB1665 + oKB1666 |
| pCG113 | pBait-CI-*Cje*-KhpB | Encodes residues 1-236 of λCI fused via three alanine residues to full-length *Campylobacter jejuni* KhpB; confers CamR | Backbone: pKB817<br>insert: oSJ5 + oSJ6 |
| pCH1 (Addgene #174663) | pBait-1xMS2$^{hp}$-empty | pCDF-pBAD-1xMS2hp. Encodes a single MS2hp under the control of an arabinose-inducible promoter, followed by XmaI and HindIII sites. CloDF13 origin of replication; confers SpecR | (Pandey et al. 2020) |
| pCH6 (Addgene #174662) | pBait-1xMS2$^{hp}$-ChiX | *E. coli chiX* inserted between XmaI/HindIII sites in pCH1; sRNA encodes its own terminator; confers SpecR | (Pandey et al. 2020) |
| pHL18 | pBait-1xMS2$^{hp}$-*sodB* | *E. coli sodB* inserted between XmaI/HindIII sites in pCH1; sRNA encodes its own terminator; confers SpecR | Backbone: pCH1<br>Insert: oHL42+43 |

| | | | |
|---|---|---|---|
| pKB816 | pBait-CI-Hfq | Encodes residues 1-236 of λCI fused via three alanine residues to *E. coli hfq*; confers CamR | (Berry and Hochschild 2018) |
| pKB817 | pPrey-α-Hfq | Encodes residues 1-248 of the alpha subunit of RNA polymerase fused via three alanine residues to full-length E. coli *hfq*; confers AmpR | (Berry and Hochschild 2018) |
| pKB845 | pBait-2xMS2$^{hp}$-XmaI-HindIII | Two MS2 RNA hairpins (2xMS2hp) and an XmaI site inserted into pKB822 CDF origin vector between BamHI and HindIII sites; confers SpecR | (Berry and Hochschild 2018) |
| pKB1288 | pPrey-α-linker-*Cje*-KhpB-KH | Encodes residues 1-248 of the alpha subunit of RNA polymerase fused via a flexible $(GGGGS)_3$ linker and three upstream alanine residues to KH domain of *Campylobacter jejuni* KhpB; confers AmpR | Backbone: pKN14<br>insert: oKB1659 + oKB1660 |
| pKB1289 | pPrey-α-linker-*Hpy*-KhpB-KH | Encodes residues 1-248 of the alpha subunit of RNA polymerase fused via a flexible $(GGGGS)_3$ linker and three upstream alanine residues to KH domain of *Helicobacter pylori* KhpB; confers AmpR | Backbone: pKN14<br>insert: oKB1661 + oKB1662 |
| pKB1293 | pBait-CI-linker-*Cje*-KhpB-KH | Encodes residues 1-236 of λCI fused via a flexible $(GGGGS)_3$ linker and three upstream alanine residues to KH domain of *Campylobacter jejuni* KhpB; confers CamR | Backbone: pKN15<br>insert: oKB1659 + oKB1660 |
| pKB1294 | pBait-CI-linker-*Hpy*-KhpB-KH | Encodes residues 1-236 of λCI fused via a flexible $(GGGGS)_3$ linker and three upstream alanine residues to KH domain of *Helicobacter pylori* KhpB; confers CamR | Backbone: pKN15<br>insert: oKB1661 + oKB1662 |
| pKN2 | pPrey-α-*Cdiff*-KhpA | Encodes residues 1-248 of the alpha subunit of RNA polymerase fused via three alanine residues to full-length *Clostridioides difficile* KhpA; confers AmpR | Backbone: pKB817<br>insert: oKN3 + oKN4 |
| pKN4 | pPrey-α-*Cdiff*-KhpB | Encodes residues 1-248 of the alpha subunit of RNA polymerase fused via three alanine residues to full-length *Clostridioides difficile* KhpB; confers AmpR | Backbone: pKB817<br>insert: oKN7 + oKN8 |
| pKN6 | pBait-CI-*Cdiff*-KhpA | Encodes residues 1-236 of λCI fused via three alanine residues to full-length *Clostridioides difficile* KhpA; confers CamR | Backbone: pKB816<br>insert: oKN3 + oKN4 |

| | | | |
|---|---|---|---|
| pKN8 | pBait-CI-*Cdiff*-KhpB | Encodes residues 1-236 of λCI fused via three alanine residues to full-length *Clostridioides difficile* KhpB; confers CamR | Backbone: pKB816 insert: oKN7 + oKN8 |
| pKN14 | pPrey-α-linker-*Cdiff*-KhpA | Encodes residues 1-248 of the alpha subunit of RNA polymerase fused via a flexible $(GGGGS)_3$ linker and three upstream alanine residues to full-length *Clostridioides difficile* KhpA; confers AmpR | Q5 on pKN2 using oKN16 + oKN17 |
| pKN15 | pBait-CI-linker-*Cdiff*-KhpA | Encodes residues 1-236 of λCI fused via a flexible $(GGGGS)_3$ linker and three upstream alanine residues to full-length *Clostridioides difficile* KhpA; confers CamR | Q5 on pKN6 using oKN15 + oKN16 |
| pKN16 | pBait-CI-linker-*Cje*-KhpA | Encodes residues 1-236 of λCI fused via a flexible $(GGGGS)_3$ linker and three upstream alanine residues to full-length *Campylobacter jejuni* KhpA; confers CamR | Backbone: pKN15 insert: oSJ1 + oSJ2 |
| pKN17 | pBait-CI-linker-*Hpy*-KhpA | Encodes residues 1-236 of λCI fused via a flexible $(GGGGS)_3$ linker and three upstream alanine residues to full-length *Helicobacter pylori* KhpA; confers CamR | Backbone: pKN15 insert: oSJ4 + oSJ9 |
| pKN21 | pBait-5GC[1xMS2$^{hp}$]-*Cdiff* nc077 | *Clostridioides difficile nc077* inserted between XmaI/HindIII sites in pSS1; sRNA encodes its own terminator; confers SpecR | Backbone: pLN28 insert: oKN20 + oKN21 |
| pKN23 | pBait-5GC[1xMS2$^{hp}$] -*Cdiff* nc013 | *Clostridioides difficile nc013* inserted between XmaI/HindIII sites in pSS1; sRNA encodes its own terminator; confers SpecR | Backbone: pLN28 insert: oKN22 + oKN24 |
| pKN24 | pBait-5GC[1xMS2$^{hp}$]-*Cdiff* nc070 | *Clostridioides difficile nc070* inserted between XmaI/HindIII sites in pSS1; sRNA encodes its own terminator; confers SpecR | Backbone: pLN28 insert: oKN25 + oKN26 |
| pKN25 | pBait-5GC[1xMS2$^{hp}$]-*Cdiff* nc008 | *Clostridioides difficile nc008* inserted between XmaI/HindIII sites in pSS1; sRNA encodes its own terminator; confers SpecR | Backbone: pLN28 insert: oKN27 + oKN28 |
| pKN26 | pBait-5GC[1xMS2$^{hp}$]-*Cdiff* nc088 | *Clostridioides difficile nc088* inserted between XmaI/HindIII sites in pSS1; sRNA encodes its own terminator; confers SpecR | Backbone: pLN28 insert: oKN29 + oKN30 |
| pKN27 | pBait-5GC[1xMS2$^{hp}$]-*Cdiff* nc083 | *Clostridioides difficile nc083* inserted between XmaI/HindIII sites in pSS1; sRNA encodes its own terminator; confers SpecR | Backbone: pLN28 insert: oKN31 + oKN32 |

| | | | |
|---|---|---|---|
| pKN29 | pPrey-α-linker-*Cje*-KhpA | Encodes residues 1-248 of the alpha subunit of RNA polymerase fused via a flexible $(GGGGS)_3$ linker and three upstream alanine residues to full-length *Campylobacter jejuni* KhpA; confers AmpR | Backbone: pKN14<br>insert: oSJ1 + oSJ2 |
| pKN30 | pPrey-α-linker-*Hpy*-KhpA | Encodes residues 1-248 of the alpha subunit of RNA polymerase fused via a flexible $(GGGGS)_3$ linker and three upstream alanine residues to full-length *Helicobacter pylori* KhpA; confers AmpR | Backbone: pKN14<br>insert: oSJ4 + oSJ9 |
| pKN32 | pPrey-α-linker-*Cdiff*-KhpA (GKQG→GDDG) | Site-directed mutant of GXXG motif residues of *Clostridioides difficile* KhpA in context of pKN14; confers AmpR | Q5 on pKN14 using oKN51 + oKN52 |
| pKN33 | pPrey-α-linker-*Cdiff*-KhpA (GKQG→AKQA) | Site-directed mutant of GXXG motif residues of *Clostridioides difficile* KhpA in context of pKN14; confers AmpR | Q5 on pKN14 using oKN52 + oKN53 |
| pKN34 | pPrey-α-linker-*Cdiff*-KhpA (GKQG→GAAG) | Site-directed mutant of GXXG motif residues of *Clostridioides difficile* KhpA in context of pKN14; confers AmpR | Q5 on pKN14 using oKN52 + oKN54 |
| pKN35 | pPrey-α-linker-*Cdiff*-KhpA (GKQG→GAQG) | Site-directed mutant of GXXG motif residues of *Clostridioides difficile* KhpA in context of pKN14; confers AmpR | Q5 on pKN14 using oKN52 + oKN55 |
| pKN36 | pPrey-α-linker-*Cdiff*-KhpA (GKQG→GKAG) | Site-directed mutant of GXXG motif residues of *Clostridioides difficile* KhpA in context of pKN14; confers AmpR | Q5 on pKN14 using oKN52 + oKN56 |
| pKN37 | pPrey-α-linker-*Cdiff*-KhpA (GKVIG→AKVIG) | Site-directed mutant of GXXG motif residues of *Clostridioides difficile* KhpA in context of pKN14; confers AmpR | Q5 on pKN14 using oKN57 + oKN58 |
| pLN35 | pBait-5GC[1xMS2$^{hp}$]-CyaR | *E. coli cyaR* inserted between XmaI/HindIII sites in pSS1; sRNA encodes its own terminator; confers SpecR | (Nguyen et al. 2025) |
| pLN91 | pBait-5GC[1xMS2$^{hp}$]-SgrS | *E. coli sgrS* inserted between XmaI/HindIII sites in pSS1; sRNA encodes its own terminator; confers SpecR | (Nguyen et al. 2025) |
| pNWL1 | pBait-5GC[1xMS2$^{hp}$]-CJnc140 | *Campylobacter jejuni* CJnc140 inserted between XmaI/HindIII sites in pSS1; sRNA encodes its own terminator; confers SpecR | Backbone: pLN28<br>insert: oNWL1 + oNWL2 |
| pNWL2 | pBait-1xMS2$^{hp}$-7GC[*Hpy thrC*-5′]-$T_{trpA}$ | *Helicobacter pylori* 5′UTR of *thrC* inserted between XmaI/HindIII sites in pSS2; | Backbone: pLN43<br>insert: oNWL3 + |

| | | | |
|---|---|---|---|
| | | confers SpecR | oNWL4 |
| pNWL5 | pBait-1xMS2$^{hp}$-7GC [Cj*0416*-5′]-$T_{trpA}$ | *Campylobacter jejuni* 5′UTR of *Cj0416* inserted between XmaI/HindIII sites in pSS2; confers SpecR | Backbone: pLN43 insert: oNWL9 + oNWL10 |
| pNWL9 | pBait-1xMS2$^{hp}$-7GC[*Hpy flgM*]-$T_{trpA}$ | *Helicobacter pylori* 5′UTR of *flgM* inserted between XmaI/HindIII sites in pSS2; confers SpecR | Backbone: pLN43 insert: oNWL16 + oKN35 |
| pNWL10 | pBait-1xMS2$^{hp}$-7GC[*Cje flgM*]-$T_{trpA}$ | *Campylobacter jejuni* 5′UTR of *flgM* inserted between XmaI/HindIII sites in pSS2; confers SpecR | Backbone: pLN43 insert: oNWL5 + oKN36 |
| pNWL12 | pBait-1xMS2$^{hp}$-7GC[*Hpy hopH*]-$T_{trpA}$ | *Helicobacter pylori* 5′UTR of *hopH* inserted between XmaI/HindIII sites in pSS2; confers SpecR | Backbone: pLN43 insert: oNWL24 + oNWL25 |
| pNWL13 | pBait-1xMS2$^{hp}$-7GC[*Cje flhB*]-$T_{trpA}$ | *Campylobacter jejuni* 5′UTR of *flhB* inserted between XmaI/HindIII sites in pSS2; confers SpecR | Backbone: pLN43 insert: oKN38 + oNWL29 |
| pNWL14 | pBait-1xMS2$^{hp}$-7GC[*Hpy cagG*]-$T_{trpA}$ | *Helicobacter pylori* 5′UTR of *cagG* inserted between XmaI/HindIII sites in pSS2; confers SpecR | Backbone: pLN43 insert: oKN39 + oKN40 |
| pNWL15 | pBait-1xMS2$^{hp}$-7GC [Cj*0604*]-$T_{trpA}$ | *Campylobacter jejuni* 5′UTR of *Cj0604* inserted between XmaI/HindIII sites in pSS2; confers SpecR | Backbone: pLN43 insert: oKN41 + oKN42 |
| pNWL17 | pBait-1xMS2$^{hp}$-7GC[*Cje ksgA*]-$T_{trpA}$ | *Campylobacter jejuni* 5′UTR of *ksgA* inserted between XmaI/HindIII sites in pSS2; confers SpecR | Backbone: pLN43 insert: oKN45 + oKN46 |
| pNWL18 | pBait-1xMS2$^{hp}$-7GC[*Hpy cagB*-5′]-$T_{trpA}$ | *Helicobacter pylori* 5′UTR of *cagB* inserted between XmaI/HindIII sites in pSS2; confers SpecR | Backbone: pLN43 insert: oKN47 + oKN48 |
| pNWL20 | pBait-1xMS2$^{hp}$-7GC[*Cje capA*-5′]-$T_{trpA}$ | *Campylobacter jejuni* 5′UTR of *capA* inserted between XmaI/HindIII sites in pSS2; confers SpecR | Backbone: pLN43 insert: oKN49 + oKN50 |
| pNWL21 | pPrey-α-linker-*Cje*-KhpA (GKNG→GDDG) | Site-directed mutant of GXXG motif residues of *Campylobacter jejuni* KhpA in context of pKN29; confers AmpR | Q5 on pKN29 using oNWL80 + oNWL81 |
| pNWL22 | pPrey-α-linker-*Cje*-KhpA (GKNG→AKNA) | Site-directed mutant of GXXG motif residues of *Campylobacter jejuni* KhpA in context of pKN29; confers AmpR | Q5 on pKN29 using oNWL52 + oNWL53 |

| | | | |
|---|---|---|---|
| pNWL23 | pPrey-α-linker-*Cje*-KhpA (GKNG→GKEG) | Site-directed mutant of GXXG motif residues of *Campylobacter jejuni* KhpA in context of pKN29; confers AmpR | Q5 on pKN29 using oNWL54 + oNWL55 |
| pNWL24 | pPrey-α-linker-*Cje*-KhpA (GKNG→GKQG) | Site-directed mutant of GXXG motif residues of *Campylobacter jejuni* KhpA in context of pKN29; confers AmpR | Q5 on pKN29 using oNWL55 + oNWL56 |
| pNWL25 | pPrey-α-linker-*Hpy*-KhpA (GKEG →GKNG) | Site-directed mutant of GXXG motif residues of *Helicobacter pylori* KhpA in context of pKN30; confers AmpR | Q5 on pKN30 using oNWL57 + oNWL58 |
| pNWL26 | pPrey-α-linker-*Hpy*-KhpA (GKEG →GKQG) | Site-directed mutant of GXXG motif residues of *Helicobacter pylori* KhpA in context of pKN30; confers AmpR | Q5 on pKN30 using oNWL58 + oNWL59 |
| pNWL28 | pPrey-α-linker-*Cje*-KhpA (GKLIG→AKLIA) | Site-directed mutant of GXXG motif residues of *Campylobacter jejuni* KhpA in context of pKN29; confers AmpR | Q5 on pKN29 using oNWL62 + oNWL63 |
| pNWL29 | pPrey-α-linker-*Cje*-KhpA (GKLIG→GKLAG) | Site-directed mutant of GXXG motif residues of *Campylobacter jejuni* KhpA in context of pKN29; confers AmpR | Q5 on pKN29 using oNWL64 + oNWL65 |
| pNWL30 | pPrey-α-linker-*Cje*-KhpA (GKLIG →GHVIG) | Site-directed mutant of GXXG motif residues of *Campylobacter jejuni* KhpA in context of pKN29; confers AmpR | Q5 on pKN29 using oNWL66 + oNWL69 |
| pNWL31 | pPrey-α-linker-*Cje*-KhpA (GKLIGKNG → GHVIGKEG) | Site-directed mutant of GXXG motif residues of *Campylobacter jejuni* KhpA in context of pKN29; confers AmpR | Q5 on pKN29 using oNWL68 + oNWL69 |
| pNWL32 | pPrey-α-linker-*Hpy*-KhpA (GHVIG →GKLIG) | Site-directed mutant of GXXG motif residues of *Helicobacter pylori* KhpA in context of pKN30; confers AmpR | Q5 on pKN30 using oNWL70 + oNWL73 |
| pNWL33 | pPrey-α-linker-*Hpy*-KhpA (GHVIGKEG → GKLIGKNG) | Site-directed mutant of GXXG motif residues of *Helicobacter pylori* KhpA in context of pKN30; confers AmpR | Q5 on pKN30 using oNWL72 + oNWL73 |
| pNWL34 | pPrey-α-linker-*Cdiff*-KhpA (GKVIG→GHVIG) | Site-directed mutant of GXXG motif residues of *Clostridioides difficile* KhpA in context of pKN14; confers AmpR | Q5 on pKN14 using oNWL74 + oNWL77 |
| pNWL35 | pPrey-α-linker-*Cdiff*-KhpA (GKQG→GKEG) | Site-directed mutant of GXXG motif residues of *Clostridioides difficile* KhpA in context of pKN14; confers AmpR | Q5 on pKN14 using oNWL75 + oNWL76 |

| | | | |
|---|---|---|---|
| pNWL36 | pPrey-α-linker-*Cdiff*-KhpA (GKVIGKQG→GHVIGKEG) | Site-directed mutant of GXXG motif residues of *Clostridioides difficile* KhpA in context of pKN14; confers AmpR | Q5 on pKN14 using oNWL76 + oNWL77 |
| pNWL37 | pPrey-α-linker-*Hpy*-KhpA (GHVIG →GKVIG) | Site-directed mutant of GXXG motif residues of *Helicobacter pylori* KhpA in context of pKN30; confers AmpR | Q5 on pKN30 using oNWL70 + oNWL79 |
| pNWL38 | pPrey-α-linker-*Hpy*-KhpA (GHVIGKEG →GKVIGKQG) | Site-directed mutant of GXXG motif residues of *Helicobacter pylori* KhpA in context of pKN30; confers AmpR | Q5 on pKN30 using oNWL78 + oNWL79 |
| pSJ1 | pPrey-α-*Cje*-KhpA | Encodes residues 1-248 of the alpha subunit of RNA polymerase fused via three alanine residues to full-length *Campylobacter jejuni* KhpA; confers AmpR | Backbone: pKB817 insert: oSJ1 + oSJ2 |
| pSJ3 | pPrey-α-*Hpy*-KhpA | Encodes residues 1-248 of the alpha subunit of RNA polymerase fused via three alanine residues to full-length *Helicobacter pylori* KhpA; confers AmpR | Backbone: pKB817 insert: oSJ4 + oSJ9 |
| pSJ5 | pBait-CI-*Cje*-KhpA | Encodes residues 1-236 of λCI fused via three alanine residues to full-length *Campylobacter jejuni* KhpA; confers CamR | Backbone: pKB816 insert: oSJ1 + oSJ2 |
| pSJ7 | pBait-CI-*Hpy*-KhpA | Encodes residues 1-236 of λCI fused via three alanine residues to full-length *Helicobacter pylori* KhpA; confers CamR | Backbone: pKB816 insert: oSJ4 + oSJ9 |
| pSP25 | pBait-1XMS2$^{hp}$-RyjB | *E. coli ryjB* (140 nts) cloned behind MS2hp in pCH1 between XmaI/HindIII sites; confers SpcR | (Pandey et al. 2020) |
| pSS1 (Addgene #222405) | pBait-5GC[1xMS2$^{hp}$] | pCDF-pBAD-5GC[1xMS2hp]. Encodes a single MS2hp flanked by 5-bp GC clamp, followed by XmaI and HindIII sites; hybrid RNA is under the control of an arabinose-inducible promoter, CloDF13 origin of replication; confers SpecR | (Nguyen et al. 2025) |
| pSS2 (Addgene #222406) | pBait-1xMS2$^{hp}$-7GC-$T_{trpA}$ | pCDF-pBAD-1xMS2hp-7GC[XmaI-HindIII]-TAA-$T_{trpA}$. Encodes a single MS2hp followed by XmaI and HindIII sites flanked by a 7-bp GC clamp, a stop codon (TAA) and the intrinsic terminator from E. coli trpA gene ($T_{trpA}$). Hybrid RNA is under the control of an arabinose-inducible promoter. | (Nguyen et al. 2025) |

| | | |
|---|---|---|
| | | CloDF13 origin of replication; confers SpecR |

**Supplementary Table S3. Oligonucleotides used in this study.** F indicates forward primer for PCR, and R indicates reverse primer. Restriction sites added through PCR are noted.

| Oligo Name | Sequence (5′ to 3′) | Details |
|---|---|---|
| oHL42 | GGCCGG CCCGGG ATACGCACAATAAGGCTATTGTACG | F XmaI Ec sodB (pHL18) |
| oHL43 | GGCCGG AAGCTT TGGTAGTGCAGGTAATTCGAATGAC | R HindIII Ec sodB (pHL18) |
| oKB1659 | GGCCGG GCGGCCGCAACTCAGGATATTTTAGATGAGATTC GTATTCAGC | F NotI Cje KhpB KH (pKB1288, pKB1293) |
| oKB1660 | CCGGCCGGATCCTCAAGCGATCTCAAGGCGCA CTAAAAG | R BamHI Cje KhpB KH (pKB1288, pKB1293) |
| oKB1661 | GGCCGGGCGGCCGCACCTAAAGAAGAATCTCAT AATGGAGACAAACTCC | F NotI Hpy KhpB KH (pKB1289, pKB1294) |
| oKB1662 | CCGGCCAGATCTTCAGGAGATTTCTAAGCGGAT GCTATAGCC | R BglII Hpy KhpB KH (pKB1289, pKB1294) |
| oKB1665 | AAAAAAGATGAGTGAGGATC | F Q5 Cje KhpB (pCG112) |
| oKB1666 | GAAAAAGTCGTTTATTACAACAAATC | R Q5 Cje KhpB (pCG112) |
| oKN3 | GGCCGGGCGGCCGCAAAAGAGCTAGTTGTAGA TATAGCTAAGGCTC | F NotI Cd KhpA (pKN2, pKN6) |
| oKN4 | CCGGCCGGATCCTTATACTATCTCAAGAGAAACT TTTATATTTTCTCTATTTGC | R BamHI Cdiff KhpA (pKN2, pKN6) |
| oKN7 | GGCCGGGCGGCCGCAAGAAAAAGCCTAAAATTA ATAAAAAGTAAGAGTAAAGAGGAAG | F NotI Cdiff KhpB (pKN4, pKN8) |
| oKN8 | CCGGCCGGATCCTTATCTTTTACACTCAATTACTA ATCTTCTATATGGATCGG | R BamHI Cdiff KhpB (pKN4, pKN8) |
| oKN15 | CTACCTCCACCTCCAGACCCGCCCCCACCGCCA AACGTCTCTTCAG | R Q5 Cdiff KhpA (pKN15) |
| oKN16 | CGGCGGTGGTGGCAGTGCGGCCGCAAAAGAG | F Q5 Cdiff KhpA (pKN14, pKN15) |
| oKN17 | CTACCTCCACCTCCAGACCCGCCCCCACCCTCT GGTTTCTCTTCTTTCAC | R Q5 Cdiff KhpA (pKN14) |
| oKN20 | GGCCGGCCCGGGAGAGAGATGAAATATATGATG AATTATAGAGAAA | F XmaI Cdiff nc077 (pKN21) |

| | | |
|---|---|---|
| oKN21 | CCGGCCAAGCTTAAACAAGAAAAAAACCTCCTTA GAGG | R HindIII Cdiff nc077 (pKN21) |
| oKN22 | GGCCGGCCCGGGATAGAAGATAGTAAAACTAGC TTAAAAACATAATATAATAC | F XmaI Cdiff nc013 (pKN23) |
| oKN24 | CATCCTCAATTGTTGCCACTATTTTAATTAAGCTT GGCCGG | R HindIII Cdiff nc013 (pKN23) |
| oKN25 | GGCCGGCCCGGGATACACGTAGATATTTATTAGT TAGATTTGAGG | F XmaI Cdiff nc070 (pKN24) |
| oKN26 | CCGGCCAAGCTTAAACAATAAGAGAGGAATTAAT CCTCTC | R HindIII Cdiff nc070 (pKN24) |
| oKN27 | GGCCGGCCCGGGACAAAATAGTATAAATAGAAG ACATGTTACTGG | F XmaI Cdiff nc008 (pKN25) |
| oKN28 | CCGGCCAAGCTTAAAAAAGGCAGAACCTATATAA ACACC | R HindIII Cdiff nc008 (pKN25) |
| oKN29 | GGCCGGCCCGGGAACCTAGTTATTCAGGAACTG AATCCAG | F XmaI Cdiff nc088 (pKN26) |
| oKN30 | CCGGCCAAGCTTAATAAAAAAATCTATACTTTTAA AAGTATAGATTCCTAAC | R HindIII Cdiff nc088 (pKN26) |
| oKN31 | GGCCGGCCCGGGATAGAAAGAACTAGCTTAAAA ACATAATATAATTACCC | F XmaI Cdiff nc083 (pKN27) |
| oKN32 | CCGGCCAAGCTTTAAAAAAATTCCCATAAACTTAT AATTTATGGGAAAAG | R HindIII Cdiff nc083 (pKN27) |
| oKN35 | CCGGCCAAGCTTAGATTGCACCGGAGCAAGAGA AGAAAC | R HindIII Hpy-flgM-5' (pNWL9) |
| oKN36 | CCGGCCAAGCTTTGCGGTATTTGCCACATAACTT TGTTGTATAG | R HindIII Cje-flgM-5' (pNWL10) |
| oKN38 | GGCCGGCCCGGGACACATATTAAAATTTTTATTTA GGCTAAAATATGGCAGG | F XmaI Cje-flhB-5' (pNWL13) |
| oKN39 | GGCCGGCCCGGGGCTAAAAATAAGAATAAAGGA ATCAAAAGTATGAAAACG | F XmaI Hpy-cagG-5' (pNWL14) |
| oKN40 | CCGGCCAAGCTTTTGGTCAGCGTTAAGCGGAGC G | R HindIII Hpy-cagG-5' (pNWL14) |
| oKN41 | GGCCGGCCCGGGAATACTTTATAAAGGTTTGAAA ATGCAAGAAAATAATTCTCC | F XmaI Cje-Cj0604-5' (pNWL15) |
| oKN42 | CCGGCCAAGCTTTTCATTCTTTTTAACAACCGCT TGCGCTTTTG | R HindIII Cje-Cj0604-5' (pNWL15) |

| | | |
|---|---|---|
| oKN45 | GGCCGGCCCGGGTTATGGTTAAAGCAAAAAAAC AATACGGACAAAATTTTTTAATC | F XmaI Cje-ksgA-5' (pNWL17) |
| oKN46 | CCGGCCAAGCTTGGCTTGGATGATTTTTGCTAAT ACGC | R HindIII Cje-ksgA-5' (pNWL17) |
| oKN47 | GGCCGGCCCGGGAATTTTAACAAAGAAAAGAAA TAAGAAATGAACCTTCAAG | F XmaI Hpy-cagB-5' (pNWL18) |
| oKN48 | CCGGCCAAGCTTGATTCCAACTATATTTAAGCATT GCATTTGATTTATTC | R HindIII Hpy-cagB-5' (pNWL18) |
| oKN49 | GGCCGGCCCGGGACTATTTTCTTATTATAGTTTTA TATTATTTTTAAATATAAACAAGAAAATATAAATATA AAAG | F XmaI Cje-capA-5' (pNWL20) |
| oKN50 | CCGGCCAAGCTTAGAATTTAAACAGGAATTTTGT ATATCTTTTGTGTATGTTTTAG | R HindIII Cje-capA-5' (pNWL20) |
| oKN51 | GGGGATGACGGAAGAATAGCTAAAGCTATAAGAA C | F Q5 Cdiff KhpA (pKN32) |
| oKN52 | AATAACCTTACCCATATCCTCTTGAGCAAC | R Q5 Cdiff KhpA (pKN32, pKN33, pKN34, pKN35, pKN36) |
| oKN53 | GCGAAGCAGGCCAGAATAGCTAAAGCTATAAGA AC | F Q5 Cdiff KhpA (pKN33) |
| oKN54 | GGGGCGGCCGGAAGAATAGCTAAAGCTATAAGA AC | F Q5 Cdiff KhpA (pKN34) |
| oKN55 | GGGGCGCAGGGAAGAATAGCTAAAGCTATAAGA AC | F Q5 Cdiff KhpA (pKN35) |
| oKN56 | GGGAAGGCCGGAAGAATAGCTAAAGCTATAAGA AC | F Q5 Cdiff KhpA (pKN36) |
| oKN57 | GGGAAGCAGGGAAGAATAGCTAAAGCTATAAGA AC | F Q5 Cdiff KhpA (pKN37) |
| oKN58 | AATAACCTTCGCCATATCCTCTTGAGCAAC | R Q5 Cdiff KhpA (pKN37) |
| oNWL1 | GGCCGGCCCGGGGAACCGAAAAACATTCATAAG AAAAACTCC | F XmaI CJnc140 (pNWL1) |
| oNWL2 | CCGGCCAAGCTTAAAAAAAGCCTAGCTAAAGGG ATTTAAGC | R HindIII CJnc140 (pNWL1) |
| oNWL3 | GGCCGGCCCGGGATAAATCAAAAATTAAACTGG ATTAAAAAGGCGATC | F XmaI Hpy-thrC-5′ (pNWL2) |
| oNWL4 | CCGGCCAAGCTTAAGAGCGAATGCAACCTTAAAT | R HindIII Hpy-thrC-5′ |

| | | |
|---|---|---|
| | GG | (pNWL2) |
| oNWL5 | GGCCGGCCCGGGGAGAGGATTATAACTAAGATC AAGGAGGCAG | F XmaI Cje-flgM-5' (pNWL10) |
| oNWL9 | GGCCGGCCCGGGATATCAAGGAAGGCAAAAATG ATTAATGG | F XmaI Cj0416-5' (pNWL5) |
| oNWL10 | CCGGCCAAGCTTTACATTTGCCTTTTTGTATAATT GTAGTTTG | R HindIII Cj0416-5' (pNWL5) |
| oNWL13 | CGGCAGATCTTCATTCATTGTTAAAGTCGTTGAT GACG | R BglII Hpy KhpB (pCG110, pCG111) |
| oNWL16 | GGCCGGCCCGGGCAATGGAATGAATATCAAATTA AAGGATTTTACAATG | F XmaI Hpy-flgM-5' (pNWL9) |
| oNWL24 | TAACCGGGCCAGCCCGCC | F XmaI Hpy-hopH-5' (pNWL12) |
| oNWL25 | GCTTGAGAGAGAGAGAGTTAGTAAGAGAGCTTT TTTCATGGTTTTTC | R HindIII Hpy-hopH-5' (pNWL12) |
| oNWL29 | CCGGCCAAGCTTTTTTGGACGTGGGTTCTTCTG | R HindIII Cje-flhB-5' (pNWL13) |
| oNWL50 | GCTTATAGGTGATGACGGAAAAATGATCAATGC | F Q5 Cje KhpA (pNWL21) |
| oNWL51 | TTACCTGTATCAACCTTATG | R Q5 Cje KhpA (pNWL21) |
| oNWL52 | AATGCCAAAATGATCAATGCTATAAAAAC | F Q5 Cje KhpA (pNWL22) |
| oNWL53 | TTTCGCTATAAGCTTACCTGTATCAAC | R Q5 Cje KhpA (pNWL22) |
| oNWL54 | TATAGGTAAAGAAGGAAAAATGATCAATGC | F Q5 Cje KhpA (pNWL23) |
| oNWL55 | AGCTTACCTGTATCAACC | R Q5 Cje KhpA (pNWL23, pNWL24) |
| oNWL56 | TATAGGTAAACAGGGAAAAATGATCAATG | F Q5 Cje KhpA (pNWL24) |
| oNWL57 | GATTGGTAAAAACGGCAAAATGGTGAG | F Q5 Hpy KhpA (pNWL25) |
| oNWL58 | ACATGCCCCATGTCTGAT | R Q5 Hpy KhpA (pNWL25, pNWL26) |
| oNWL59 | GATTGGTAAACAGGGCAAAATGGTG | F Q5 Hpy KhpA (pNWL26) |

| | | |
|---|---|---|
| oNWL62 | TATAGCCAAAAATGGAAAAATGATCAATGC | F Q5 Cje KhpA (pNWL28) |
| oNWL63 | AGCTTCGCTGTATCAACCTTATGTGC | R Q5 Cje KhpA (pNWL28) |
| oNWL64 | AGGTAAGCTTGCGGGTAAAAATGGAAAAATG | F Q5 Cje KhpA (pNWL29) |
| oNWL65 | GTATCAACCTTATGTGCAAAAAG | R Q5 Cje KhpA (pNWL29) |
| oNWL66 | GGTAAAAATGGAAAAATGATCAATGCT | F Q5 Cje KhpA (pNWL30) |
| oNWL68 | GGTAAAGAAGGAAAAATGATCAATGCTATA | F Q5 Cje KhpA (pNWL31) |
| oNWL69 | TATCACATGACCTGTATCAACCTT | R Q5 Cje KhpA (pNWL30, pNWL31) |
| oNWL70 | GGTAAAGAGGGCAAAATGGT | F Q5 Hpy KhpA (pNWL32, pNWL37) |
| oNWL72 | GGTAAAAACGGCAAAATGGTG | F Q5 Hpy KhpA (pNWL33) |
| oNWL73 | AATCAGTTTCCCCATGTCTGA | R Q5 Hpy KhpA (pNWL32, pNWL33) |
| oNWL74 | GGGAAGCAGGGAAGAATAG | F Q5 Hpy KhpA (pNWL34) |
| oNWL75 | AATAACCTTACCCATATCCTCTTGAG | F Q5 Hpy KhpA (pNWL35, pNWL36) |
| oNWL76 | GGGAAGGAAGGAAGAATAGCT | R Q5 Hpy KhpA (pNWL35, pNWL36) |
| oNWL77 | AATAACATGACCCATATCCTCTTG | R Q5 Hpy KhpA (pNWL34) |
| oNWL78 | GGTAAACAGGGCAAAATGGT | F Q5 Hpy KhpA (pNWL38) |
| oNWL79 | AATCACTTTCCCCATGTCTGA | R Q5 Hpy KhpA (pNWL37, pNWL38) |
| oSJ1 | GCCGGCGGCCGCAGTAGAGAATTTTCTTAGAGA ATACGC | F NotI Cje KhpA (pSJ1, pSJ5, pKN16, pKN29) |
| oSJ2 | CCGGCCGGATCCTCACTCAAGTGCTTTTACCGT TACTC | R BamHI Cje KhpA (pSJ1, pSJ5, pKN16, pKN29) |

| | | |
|---|---|---|
| oSJ4 | CCGGCCGGATCCTCAAGGCGTCTGATCGCCTAAAAC | R BamHI Hpy KhpA (pSJ3, pSJ7, pKN17, pKN30) |
| oSJ5 | GCCGGCGGCCGCAAAAATCGAAGCCATTGATCTTCAAAG | F NotI Cje KhpB (pCG113) |
| oSJ6 | CGGCGGATCCTCACTCATCTTTTTTGAAAAAGTCG | R BamHI Cje KhpB (pCG113) |
| oSJ7 | GCCGGCGGCCGCACAAAATCCTATTGAAATCAAAGCCAAAAC | F NotI Hpy KhpB (pCG110, pCG111) |
| oSJ9 | GGCCGGGCGGCCGCACGTGAATTAGATCCTTTGAGCACGC | F NotI Hpy KhpA (pSJ3, pSJ7, pKN17, pKN30) |

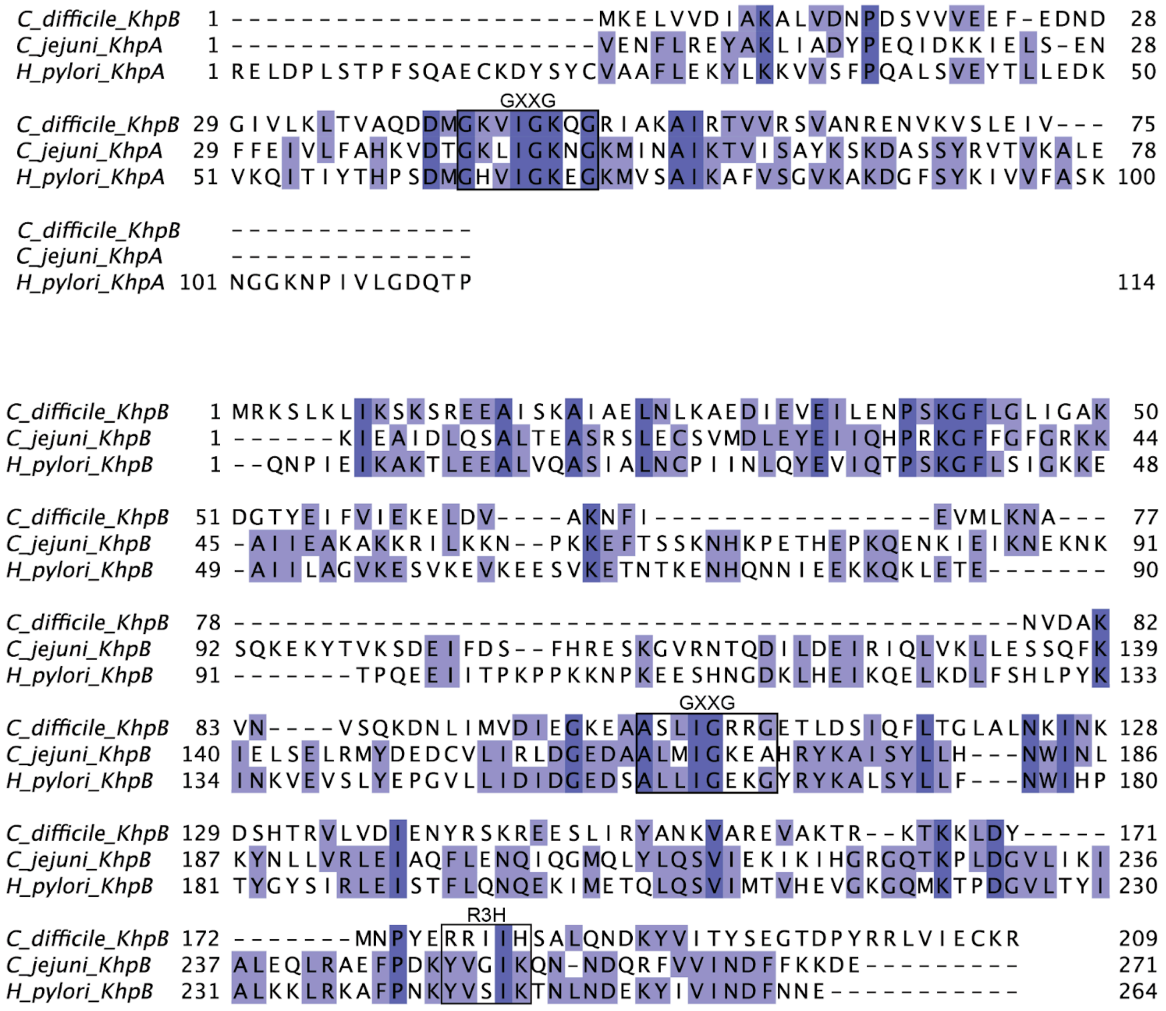

**Supplementary Figure S1**. **Sequence alignments of proteins analyzed in this study.** Alignments of KhpA (top) and KhpB (bottom) sequences were generated with Clustal Omega (Madeira et al. 2024) and visualized in Jalview (Waterhouse et al. 2009). Amino acids are colored based on degree of conservation; the GXXG and R3H motifs are boxed and labeled.

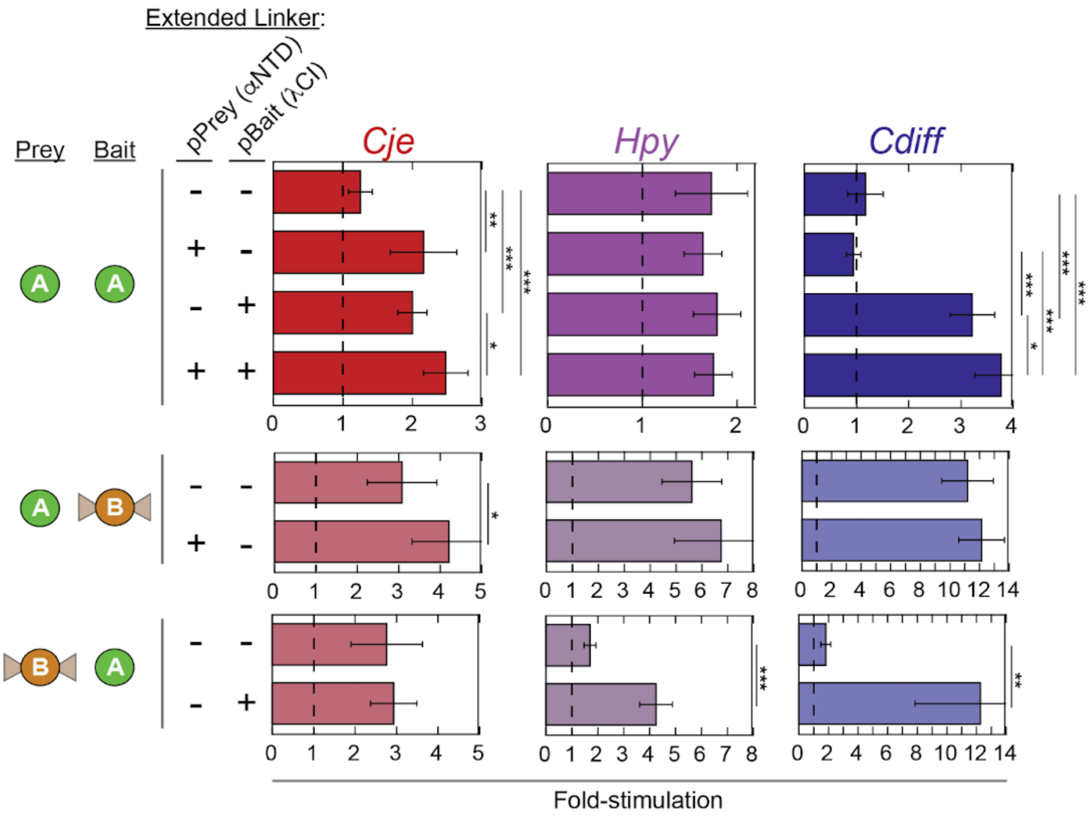

**Supplementary Figure S2. Extended linkers improve KhpA B2H interactions.** (A) Schematic comparing original and extended-linker B2H constructs. Original vectors contain a short triple-alanine (AAA) linker between the λCI or α-NTD fusion moiety and the protein of interest (Dove et al. 1997). Extended-linker constructs include a longer flexible $(GGGGS)_3$-AAA linker sequence and are indicated with an asterisk (*) in front of the schematic circle representing KhpA throughout the paper. (B) Effect of linker extension on KhpA and KhpB interactions. B2H assays were performed in KB473 reporter cells to test homodimerization of KhpA and heterodimerization of KhpA and KhpB. Transformations included combinations of pPrey and pBait constructs from *C. jejuni* (*Cje*, red; pSJ1, pKN16, pCG112, pSJ5, pKN29, pCG113), *H. pylori* (*Hpy*, purple; pKN17, pSJ3, pCG110, pKN30, pSJ7, pCG111), and *C. difficile* (*Cdiff*, blue; pKN2, pKN14, pKN4, pKN6, pKN15, pKN8) or corresponding empty vectors to establish basal transcription levels. Asterisks indicate statistically significant differences in interactions (*$P$ < 0.05, **$P$ < 0.005, ***$P$ < 0.0005) as determined by Student's t-tests.

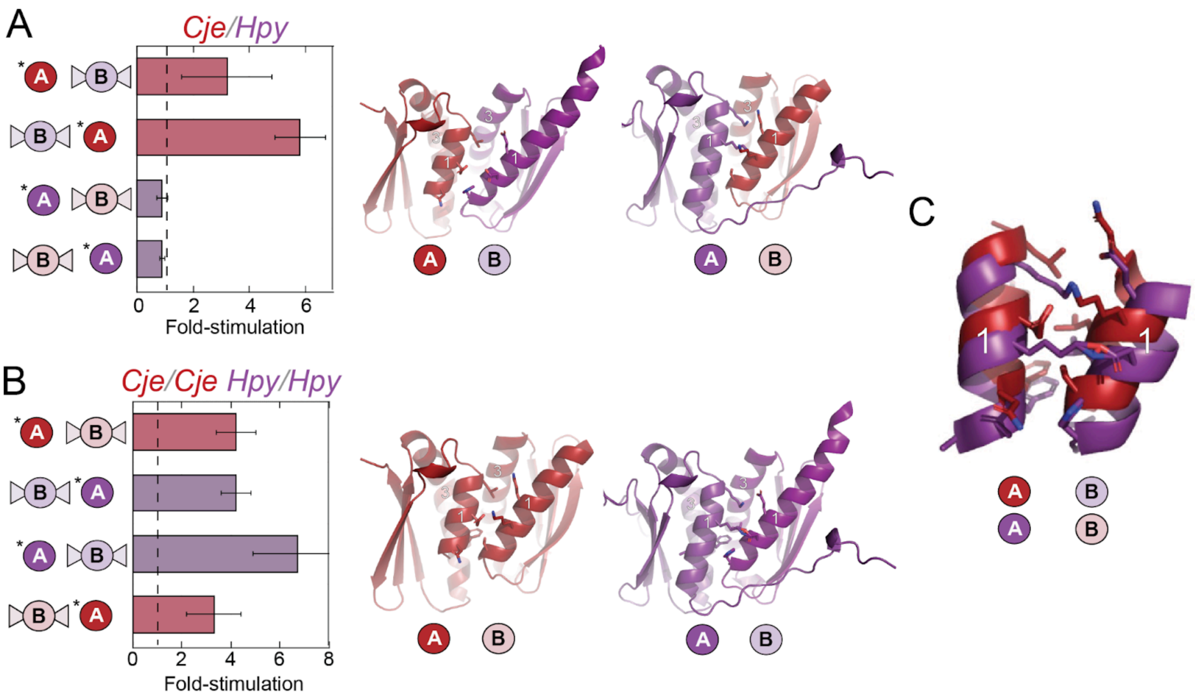

**Supplementary Figure S3. Species-specific heterodimerization of KhpA and KhpB.** Assays were performed in KB473 reporter cells as described for Figure 3 to measure protein dimerization between pPrey α-NTD fusion proteins and pBait λCI fusion proteins. (A, left) B2H data for cross-species heterodimerization interactions between *Cje* (red; pKN29, pCG112) and *Hpy* (purple; pKN30, pCG110) KhpA and KhpB in both orientations (pPrey and pBait). Assays were conducted as in Figure 3, but with cross-species combinations of *Cje* and *Hpy* constructs. (A, right) Cartoon representation of Alphafold3 predictions of structures of the KH domains of KhpA and KhpB in cross-species heterodimerization (Abramson et al. 2024). (B) Corresponding B2H data and AlphaFold predictions for the intra-species heterodimerization interactions. B2H data on the left (utilizing pKN29, pCG112, pKN30 and pCG110) are replotted from Figure 3. (C) Overlay of helix 1 of *Cje* (red) and *Hpy* (purple) from KhpA and KhpB's KH domains. Key side chains pointing to the dimerization interface are shown as sticks.

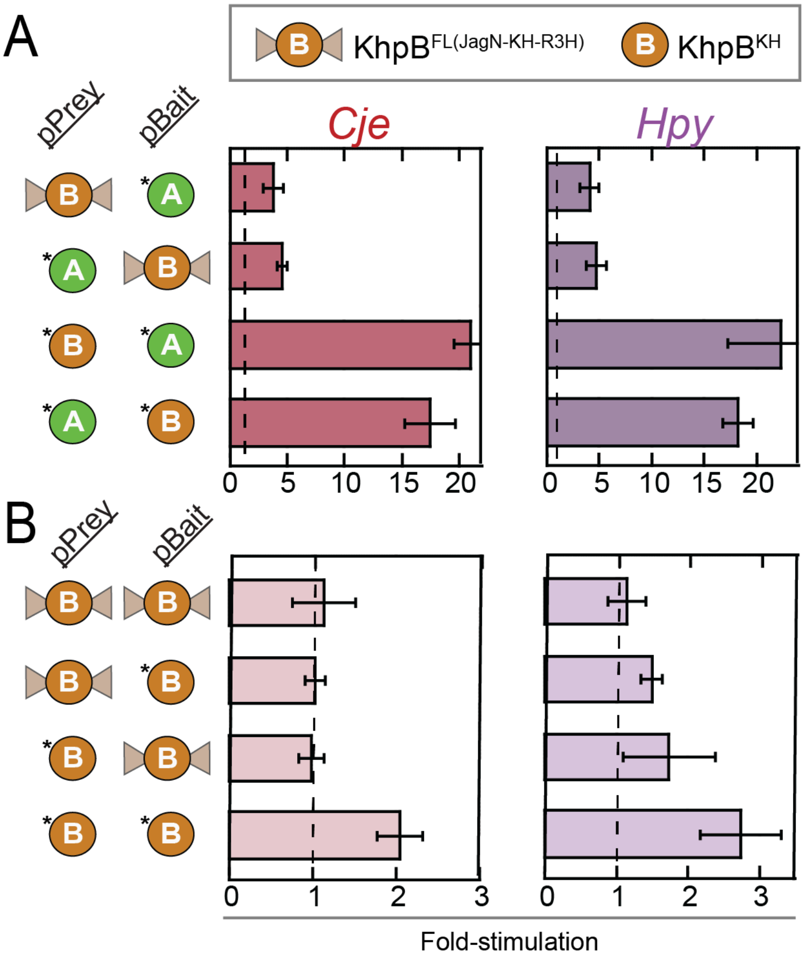

**Supplementary Figure S4. Hetero- and homo-dimerization of isolated KH domains of KhpB proteins.** Assays were performed in KB473 reporter cells as described for Figure 3 to measure protein dimerization between pPrey α-NTD fusion proteins and pBait λCI fusion proteins. Results of B2H assays detecting (A) heterodimerization of KhpA and KhpB and (B) homodimerization of KhpB using proteins from *C. jejuni* (left, red; pPrey: pKN29, pCG112 or pKB1288; pBait: pKN16, pCG113 or pKB1293) and *H. pylori* (right, purple; pPrey: pKN30, pCG110 or pKB1289; pBait: pKN17, pCG111 or pKB1294). Interactions of full-length KhpB proteins (indicated by a circle with triangles on either side) were compared to truncation constructs that contained only the KH domain of KhpB (indicated by a simple circle). All constructs containing the isolated KH domain of KhpB included an extended flexible $(GGGGS)_3$-AAA linker indicated by an asterisk (*) on the schematic diagrams.

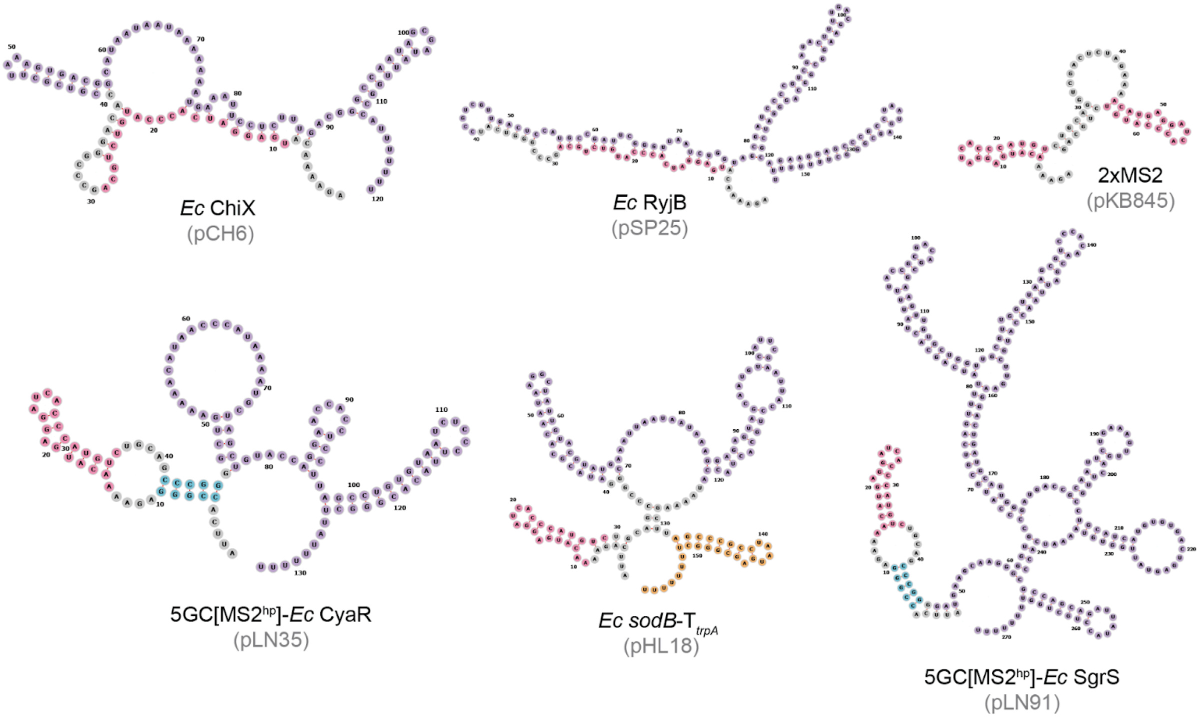

**Supplementary Figure S5. Predicted secondary structures for B3H pBait constructs containing *E. coli* RNAs.** These *E. coli* pBait-RNA constructs were used in B3H experiments shown in Figures 5, S9 and S10. RNA structure predictions and visualizations here and in subsequent figures were generated using RNAfold and *forna* (Kerpedjiev et al. 2015; Lorenz et al. 2011). Nucleotides within secondary structures are colored based on the following scheme: red = $MS2^{hp}$ moiety; blue = GC clamp; purple = inserted bait RNA; yellow = $T_{trpA}$ terminator; grey = linker region and restriction sites.

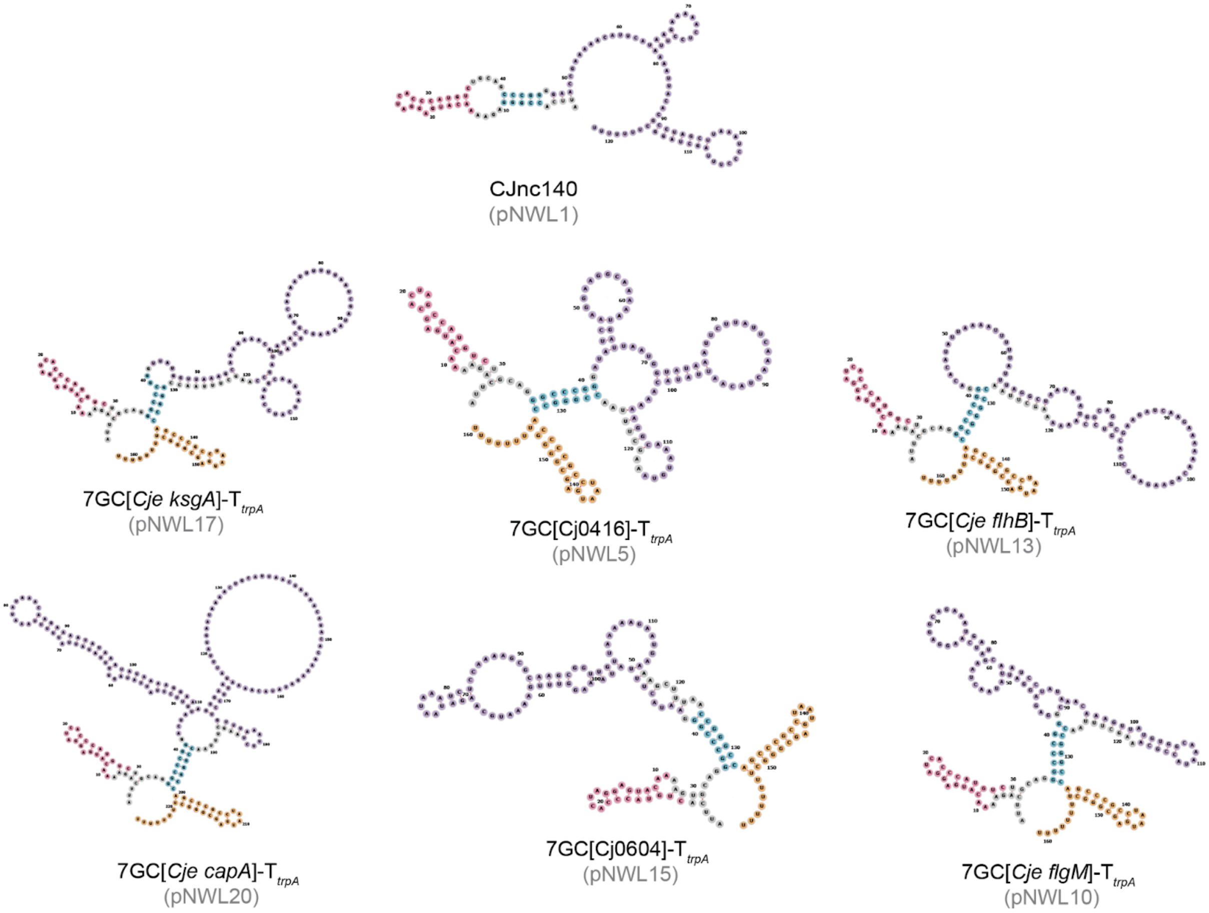

**Supplementary Figure S6. Predicted secondary structures for B3H pBait constructs containing *C. jejuni* RNAs.** These *C. jejuni* pBait-RNA constructs were used in B3H experiments shown in Figures 5, S9 and S10. Secondary structures were predicted and visualized as described for Figure S5.

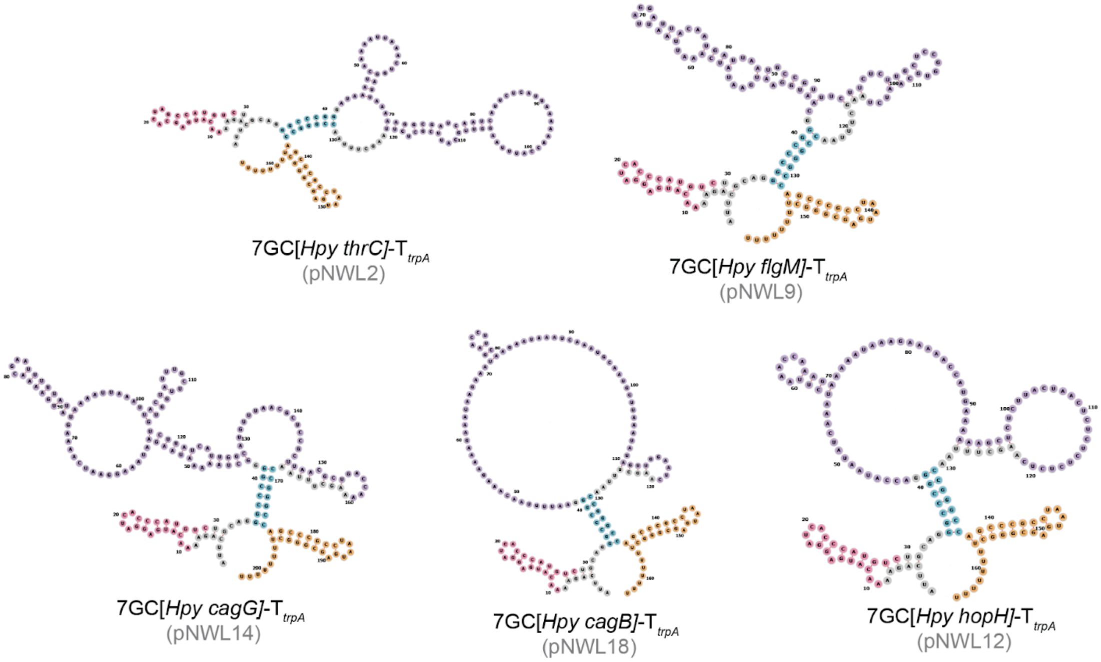

**Supplementary Figure S7. Predicted secondary structures for B3H pBait constructs containing *H. pylori* RNAs.** These *H. pylori* pBait-RNA constructs were used in B3H experiments shown in Figures 5, S9 and S10. Secondary structures were predicted and visualized as described for Figure S5.

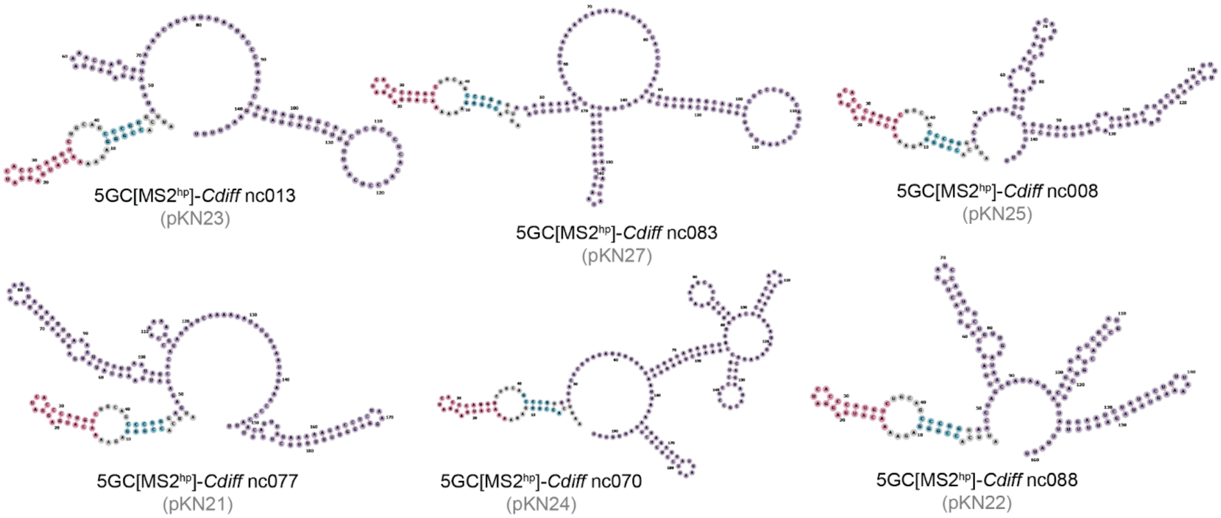

**Supplementary Figure S8. Predicted secondary structures for B3H pBait constructs containing *C. difficile* RNAs.** These *C. difficile* pBait-RNA constructs were used in B3H experiments shown in Figures 5, S9 and S10. Secondary structures were predicted and visualized as described for Figure S5.

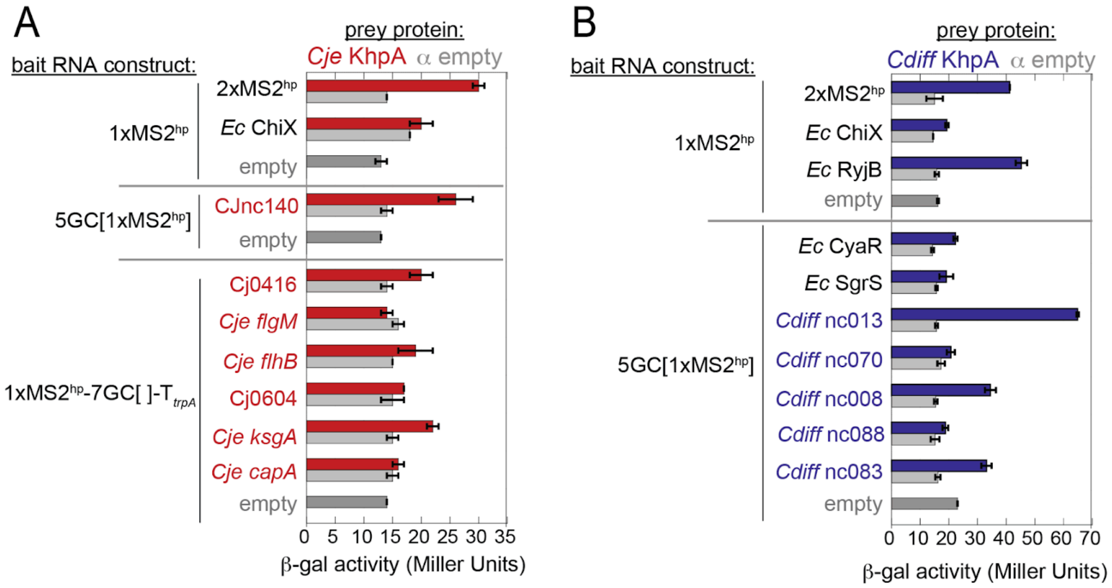

**Supplementary Figure S9. Raw β-gal data for B3H-based detection of KhpAB-RNA interactions with species-specific RNAs.** These raw β-gal activity values were measured in KB483 reporter cells to calculate the fold-stimulation data presented in Figure 5. A) *C. jejuni* and (B) *C. difficile* interactions**.** Assays utilized pPrey encoding α alone (grey bars), or α-linker-KhpA fusion proteins (pKN29 for *Cje* or pKN14 for *Cdiff*), pAdapter (pCW17), and pBait–RNA constructs encoding either the indicated empty negative-control constructs or species-specific bait RNA sequences. To calculate fold-stimulation, each experimental β-gal value was divided by the highest value from its matching negative control (containing either the α- or pBait-empty construct). Matching negative controls for pBait constructs are as follows: the $MS2^{hp}$-only moiety (pCH1) served as the control for 2xMS2hp (pKB845) and *E. coli* ChiX and RyjB (pCH6, pSP25); the construct with a 5-bp GC clamp flanking the $MS2^{hp}$ (pSS1) was used for CJnc140 (pNWL1), *E. coli* CyaR and SgrS (pLN35, pLN91), and all *Cdiff* RNAs (pKN21, pKN23–27); and the $MS2^{hp}$ moiety followed by a 7-bp GC clamp and $T_{trpA}$ (pSS2) was used for all other *Cje* RNAs (pNWL5, pNWL10, pNWL13, pNWL15, pNWL17, pNWL20).

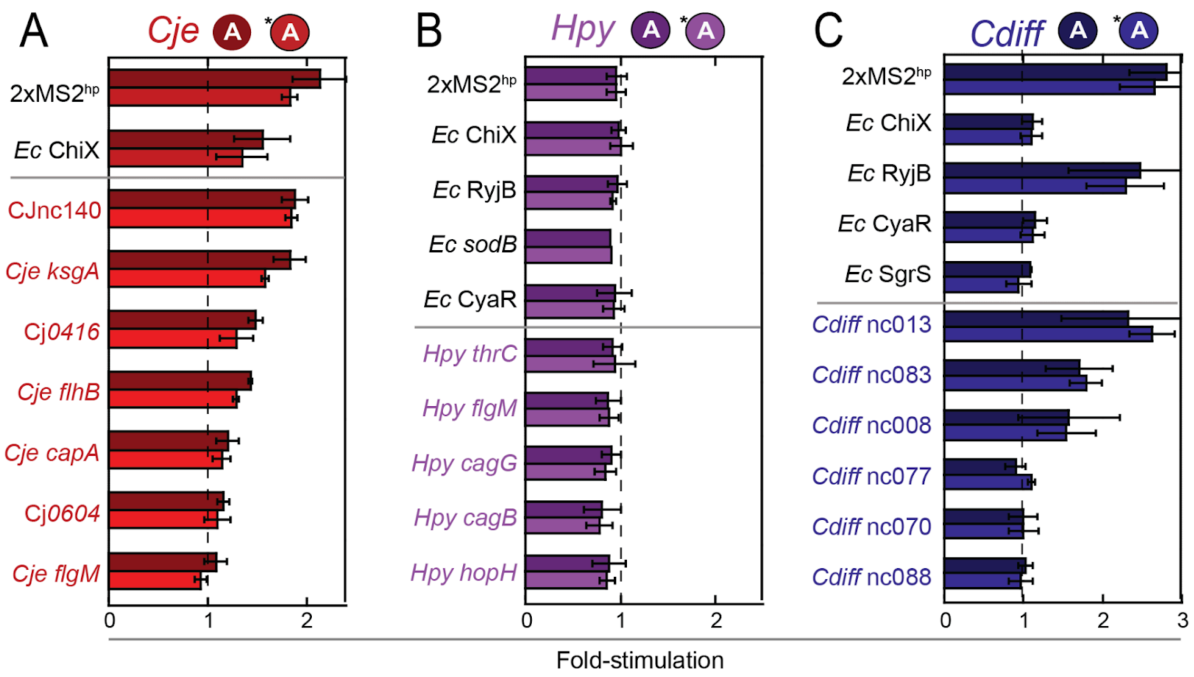

**Supplementary Figure S10. Effect of extended linker on KhpA interactions with species-specific RNAs.** B3H assays were performed in KB483 reporter cells as described for Figure 5 to measure the RNA binding of KhpA prey constructs to species-specific pBait–RNA panels from (A) *C. jejuni*, (B) *H. pylori*, and (C) *C. difficile*. Assays compared interactions of KhpA pPrey fusion proteins expressed either in the original AAA-linker constructs (darker bars; pSJ1, pSJ3, or pKN2) or in the extended $(GGGGS)_3$-AAA linker constructs (lighter bars; pKN29, pKN30, or pKN14).

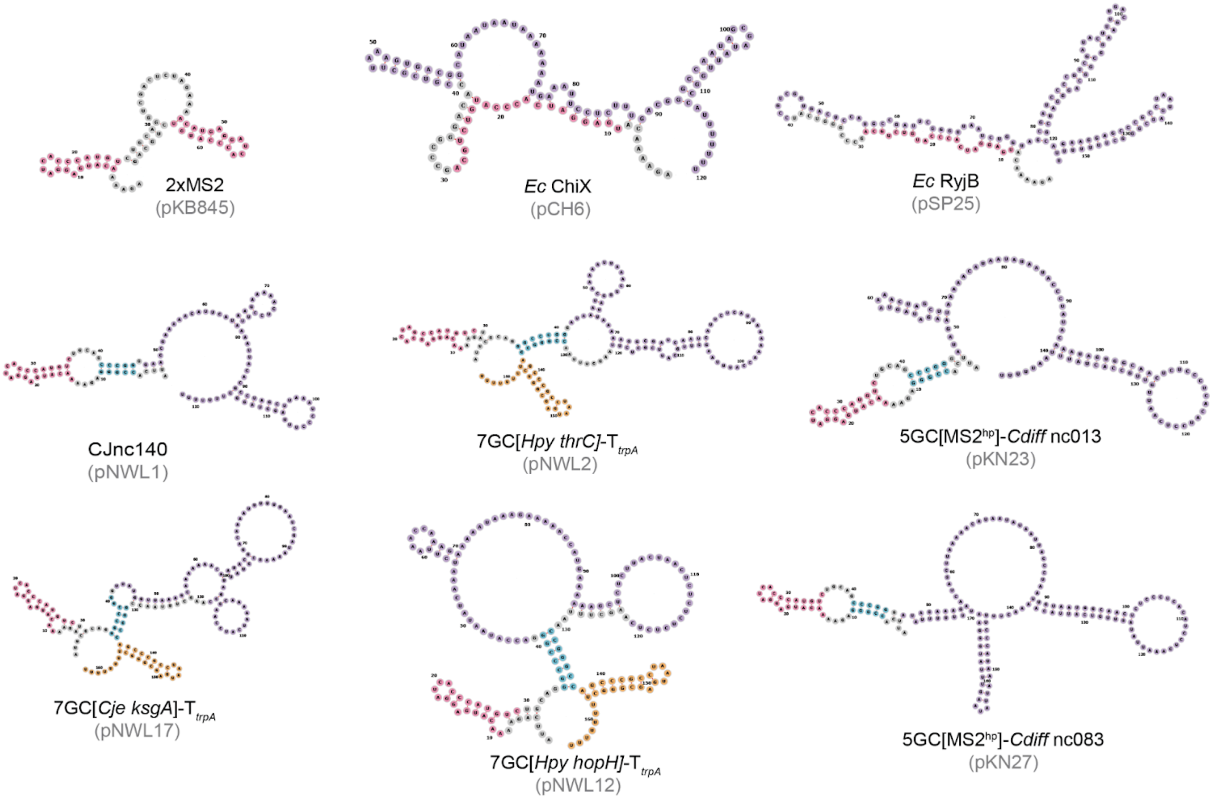

**Supplementary Figure S11. Secondary structure predictions of B3H pBait constructs selected for cross-species panel**. These bait RNA constructs were used in B3H experiments shown in Figure 6 and S12. Secondary structures were predicted and visualized as described for Figure S5.

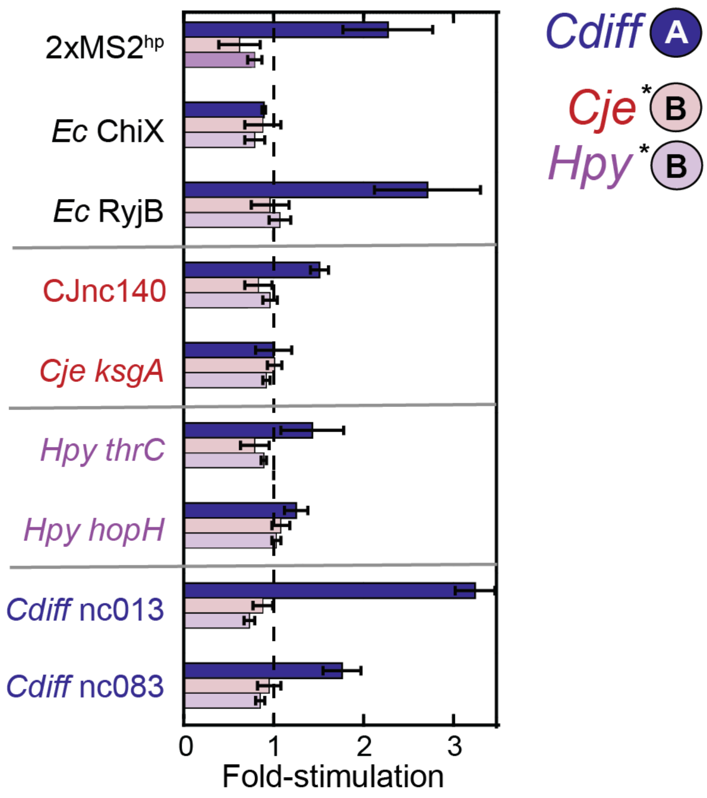

**Supplementary Figure S12. Isolated KH domains from KhpB do not demonstrate B3H interactions with RNA.** Results of B3H assays detecting RNA interactions using pPrey proteins expressing the isolated KhpB KH domains from *C. jejuni* (pKB1288; light red), *H. pylori* (pKB1289; light purple), or from *C. difficile* KhpA (pKN2; dark blue) as a side-by-side positive control for RNA interactions. Assays were conducted as in Figure 6, but with the different pPrey constructs indicated. Note that both constructs containing the isolated KH domain of KhpB included an extended $(GGGGS)_3$-AAA flexible linker.

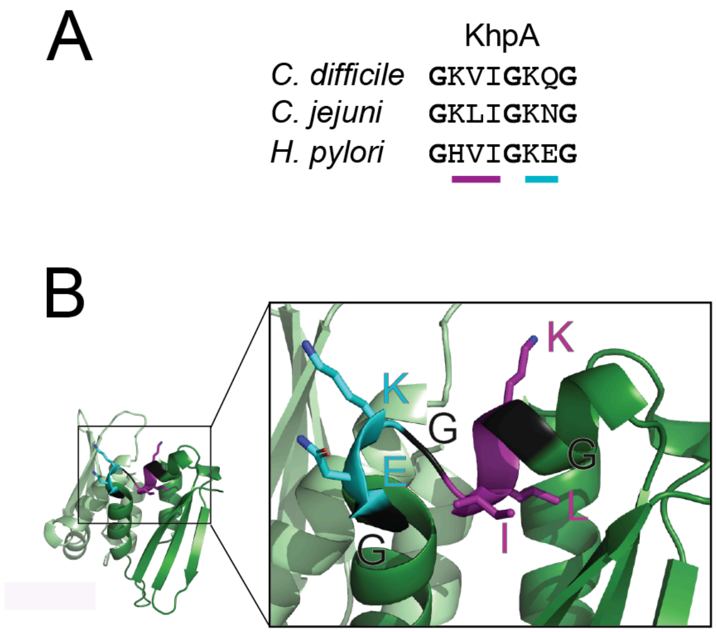

**Supplementary Figure S13. Comparison of GXXG motif residues of KhpA and KhpB proteins.** (A) GXXG motif residues found in KhpA proteins from the three species indicated. The core GXXG residues are indicated with a cyan bar, and additional “upstream” residues (GXXIG) are indicated with a purple bar. (B) Structure of GXXG motif residues within an Alphfold3 prediction of the *Cje* KhpA homodimer (Abramson et al. 2024). Conserved glycines are colored black, and additional residues are shown as sticks and colored as indicated in panel A. Two copies of this motif are present within the homodimer -- one on each KhpA monomer -- and the inset shows a close-up of the structure of one of the two motifs.

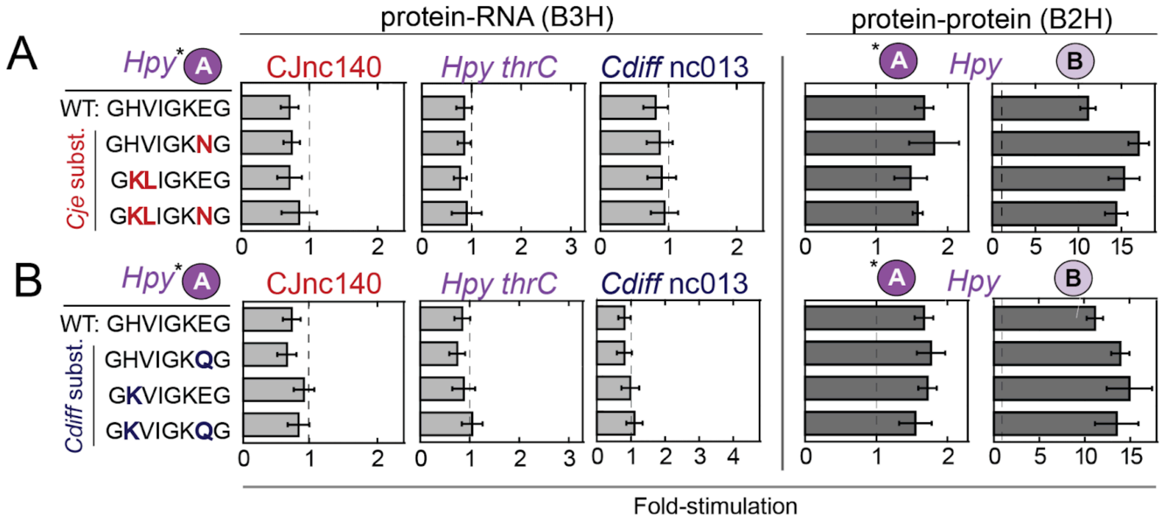

**Supplementary Figure S14. Introduction of *Cdiff* or *Cje* GXXG residues is not sufficient to restore *Hpy* KhpA-RNA interaction.** In both panels, B3H assays (left) were performed in KB483 reporter cells to measure RNA binding of KhpA pPrey constructs to pBait–RNA constructs (pNWL1, pNWL2, pKN23) or a 1xMS2$^{hp}$ negative-control construct (pCH1), while B2H assays (right) were performed in KB473 cells to measure protein dimerization of the same KhpA prey panel with pBait–protein constructs. Analysis of *Hpy* KhpA variants containing (A) *Cje*-type and (B) *Cdiff*-type GXXG residues. Assays utilized WT (pKN30) or mutant (pNWL25, pNWL26, pNWL32, pNWL33, pNWL37, and pNWL38) *Hpy* KhpA pPrey constructs. B2H assays measured dimerization with pBait constructs expressing *Hpy* KhpA (pKN17) or the isolated KH domain of *Hpy* KhpB (pKB1294). In all panels, asterisks indicate statistical significance of mutant interactions relative to WT (**P* < 0.05, ***P* < 0.005, ****P* < 0.0005) as determined by Student's t-tests.

### SUPPLEMENTARY REFERENCES

Abramson J, Adler J, Dunger J, Evans R, Green T, Pritzel A, Ronneberger O, Willmore L, Ballard AJ, Bambrick J, et al. 2024. Accurate structure prediction of biomolecular interactions with AlphaFold 3. *Nature* **630**: 493–500.

Berry KE, Hochschild A. 2018. A bacterial three-hybrid assay detects *Escherichia coli* Hfq-sRNA interactions *in vivo*. *Nucleic Acids Res* **46**: e12.

Dove SL, Joung JK, Hochschild A. 1997. Activation of prokaryotic transcription through arbitrary protein-protein contacts. *Nature* **386**: 627–630.

Kerpedjiev P, Hammer S, Hofacker IL. 2015. Forna (force-directed RNA): Simple and effective online RNA secondary structure diagrams. *Bioinformatics* **31**: 3377–3379.

Lorenz R, Bernhart SH, Höner Zu Siederdissen C, Tafer H, Flamm C, Stadler PF, Hofacker IL. 2011. ViennaRNA Package 2.0. *Algorithms Mol Biol* **6**: 26.

Madeira F, Madhusoodanan N, Lee J, Eusebi A, Niewielska A, Tivey ARN, Lopez R, Butcher S. 2024. The EMBL-EBI Job Dispatcher sequence analysis tools framework in 2024. *Nucleic Acids Res* **52**: W521–W525.

Nguyen LD, LeBlanc H, Berry KE. 2025. Improved constructs for bait RNA display in a bacterial three-hybrid assay. *Sci Rep* **15**: 3820.

Pandey S, Gravel CM, Stockert OM, Wang CD, Hegner CL, LeBlanc H, Berry KE. 2020. Genetic identification of the functional surface for RNA binding by *Escherichia coli* ProQ. *Nucleic Acids Res* **48**: 4507–4520.

Wang CD, Mansky R, Leblanc H, Gravel CM, Berry KE. 2021. Optimization of a bacterial three-hybrid assay through *in vivo* titration of an RNA–DNA adapter protein. *RNA* **27**: 513–526.

Waterhouse AM, Procter JB, Martin DMA, Clamp M, Barton GJ. 2009. Jalview Version 2—a multiple sequence alignment editor and analysis workbench. *Bioinformatics* **25**: 1189–1191.
